# Criterion for assessing accumulated neurotoxicity of alpha-synuclein oligomers in Parkinson’s disease

**DOI:** 10.1101/2024.09.23.614584

**Authors:** Andrey V. Kuznetsov

## Abstract

The paper introduces a parameter called “accumulated neurotoxicity” of α-syn oligomers, which measures the cumulative damage these toxic species inflict on neurons over time, given the years it typically takes for such damage to manifest. A threshold value for accumulated neurotoxicity is estimated, beyond which neuron death is likely. Numerical results suggest that rapid deposition of α-syn oligomers into fibrils minimizes toxicity, indicating that the formation of Lewy bodies might play a neuroprotective role. Strategies such as reducing α-syn monomer production or enhancing degradation can decrease accumulated toxicity. In contrast, slower degradation (reflected by longer half-lives of monomers and free aggregates) increases toxicity, supporting the idea that impaired protein degradation may contribute to Parkinson’s disease progression. The study also examines the sensitivity of accumulated toxicity to different model parameters.

## 1. Introduction

Parkinson’s disease (PD) is a neurodegenerative condition marked by the degeneration of dopaminergic neurons in the substantia nigra, accompanied by the formation of Lewy bodies (LBs), which are abnormal protein aggregates found within affected neurons [1-5].

Alpha-synuclein (α-syn), a key protein implicated in PD, undergoes misfolding and aggregation, resulting in the formation of oligomers that contribute to neuronal damage and toxicity [6]. Recent research suggests that LBs, traditionally viewed as toxic deposits of misfolded and aggregated α-syn within neurons, may be neuroprotective and sequester toxic α-syn oligomers [7]. This emerging perspective challenges the conventional view of LBs as purely pathological. Consequently, the relationship between LB formation and the accumulation of α-syn oligomers in the cytosol has become a critical area of investigation. Understanding the role α-syn aggregates play in toxicity and how their sequestration into LBs impacts disease progression is essential for developing therapeutic strategies targeting alpha-synucleinopathies [8].

Classical LBs [9] were traditionally understood to consist primarily of α-syn fibrils. This hypothesis was supported by extensive experimental evidence [10-12]. However, recent findings [13] reveal that LBs also contain lipid membrane fragments and altered organelles, challenging the previous notion that α-syn fibrils are the main components of LBs. This discovery has sparked ongoing debates about the role of α-syn fibrils within LBs and LB composition [12,14], suggesting that LBs may represent more complex aggregates of various intracellular materials. Classical brainstem-type LBs are characterized by a dense central core surrounded by a halo (an outer layer) of radiating filaments [11,14]. Both the core and the halo exhibit a spherical structure. It is likely that LBs initially develop from pale bodies, which are granular entities lacking a halo [14]. Over time, these pale bodies can evolve into the classical LBs with a surrounding halo [15].

To address these complexities, in ref. [16] a two-stage model of LB formation was introduced. Drawing from recent experimental findings showing that classical LBs feature a dense core surrounded by a halo of radiating filaments [12,13], the model suggests that the core of the LB develops from the aggregation of lipid membrane fragments and altered organelles. In the second stage, the surface of this core triggers the growth of α-syn fibrils, which incorporate α-syn monomers from the cytosol (Fig. 1). This model not only explains the structural organization of LBs but also shows how fibril formation restricts further core growth.

**Fig. 1.**
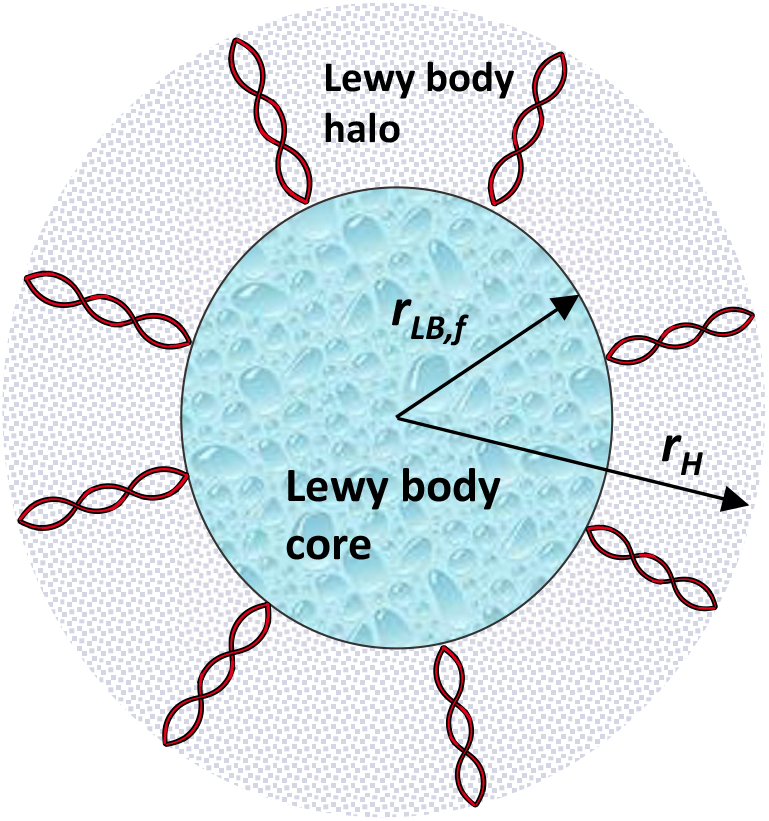
A diagram illustrating an LB, composed of two distinct regions: a central core filled with dense granular material, and a surrounding halo characterized by radiating filaments. This depiction is based on electron micrographs shown in Fig. 1 of ref. [30] and Fig. 3 of ref. [14]. *Figure generated with the aid of servier medical art, licensed under a creative commons attribution 3.0 generic license. http://Smart.servier.com*.

Developing a criterion to assess brain tissue damage at the cellular scale is important for optimizing treatments for neurodegenerative diseases [17-19]. Recent research indicates that soluble α-syn oligomers, which are intermediates in the formation of α-syn fibrils, may be more toxic than LBs [20]. Forming of these oligomeric species is a crucial step in the α-syn cascade hypothesis [21]. One potential mechanism underlying the toxicity of α-syn oligomers involves their ability to disrupt membrane integrity [22]. This study introduces a criterion for evaluating accumulated neurotoxicity (hereafter accumulated toxicity).

Additionally, it conducts a numerical analysis of α-syn aggregation, with a particular emphasis on the accumulation of toxicity over time as α-syn oligomers either persist in the cytosol or become incorporated into LBs. Building on previous research on LB growth dynamics [16,23] and recent work on the concept of accumulated toxicity of Aβ and tau oligomers [24,25], this investigation seeks to identify the key factors driving neurotoxicity in PD. It specifically examines how the sensitivity of accumulated toxicity responds to the deposition rates of α-syn oligomers into LBs, analyzing their impact on the expansion of the LB halo and overall neuronal health. It also examines how the half-lives of α-syn monomers and aggregates impact the progression of PD, highlighting the connection between impaired protein degradation pathways and disease development.

The results suggest that rapid deposition of α-syn oligomers into LBs may significantly reduce accumulated toxicity, supporting the notion that LBs could mitigate neurotoxic effects. In contrast, slow deposition allows oligomers to remain in the cytosol, where they continue to catalyze their own production and exacerbate neurotoxicity. These insights provide a framework for assessing the role of α-syn oligomers in PD and offer potential avenues for therapeutic intervention aimed at reducing their toxic effects.

## 2. Materials and models

### 2.1. Equations governing lipid membrane fragment aggregation in LB’s core

A minimal two-step Finke-Watzky (F-W) model was applied to simulate the aggregation of lipid fragments in the soma, based on approaches outlined in refs. [26,27]. This model captures polymer aggregation through two pseudoelementary processes: continuous nucleation and autocatalytic surface growth [28]. The F-W model has previously been utilized to analyze the aggregation behavior of neurological proteins [26]. The two pseudoelementary steps in the F-W model are described by the following equations:

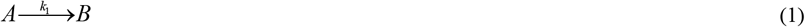

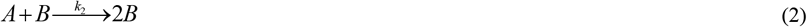

The F-W model is typically used to simulate the transformation of a given initial concentration of monomers (*A*) into misfolded or autocatalytically active aggregates (*B*) [27]. While analytical solutions for such processes are well-documented (e.g., ref. [29]), this study adapts the F-W model to a different context by simulating the scenario where monomers are continuously produced. Recent findings [13,14] indicate that the core of LBs mainly consists of lipid membrane fragments and damaged organelles, referred to here simply as membrane fragments.

The model uses time, *t*, as the sole independent variable. The dependent variables are detailed in Table S1 of the Supplemental Materials, and Table S2 summarizes the parameters used in the model, which are estimated from data reported in refs. [30-39].

The production rate of membrane fragments in the soma, *q*_*FR*_, is assumed to be constant. By considering the soma as the control volume and applying the principle of conservation of membrane fragments within this control volume, the following equation is derived:

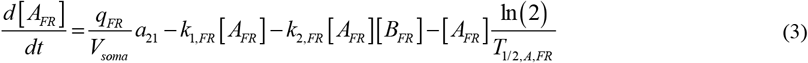

In this equation, [*A*_*FR*_] represents the concentration of membrane fragments, [*B*_*FR*_] denotes the concentration of free lipid membrane aggregates, which can form the LB core, and *a*_21_ is the conversion factor from 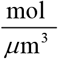 to 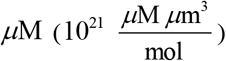. The parameters *k*_1, *FR*_ and *k*_2, *FR*_ are rate constants for the nucleation and autocatalytic growth of lipid membrane fragment aggregates, respectively. The quantity *T*_1/ 2, *A, FR*_ is the half-life of lipid membrane fragments, and *V*_*soma*_ is the volume of the soma.

The right-hand side of Eq. (3) includes several terms. The first term represents the production rate of membrane fragments. The second term models the conversion of these fragments into aggregates through nucleation, while the third term accounts for the conversion through autocatalytic growth. The final term models the degradation of membrane fragments.

By stating the conservation of membrane aggregates in the soma, the following equation is derived:

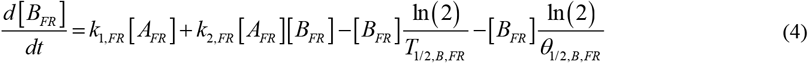

where *T*_1/ 2, *B, FR*_ represents the half-life of free aggregated lipid membrane fragments (those not yet deposited into the LB core), and *θ*_1/ 2, *B, FR*_ denotes the half-deposition time of free lipid membrane aggregates into the LB core. On the right-hand side, the first term captures the nucleation-driven rate of free aggregate production from monomers, while the second term describes the rate of free aggregate production through autocatalytic growth. These terms mirror the corresponding ones in Eq. (3) but with opposite signs, as the F-W model assumes that aggregate production equals monomer consumption. The third term models the degradation of lipid membrane aggregates, and the fourth term represents their deposition into the LB core.

The process of LB core formation from adhesive lipid fragment aggregates is modeled by an approach akin to colloidal suspension coagulation [40]. In this framework, free lipid fragment aggregates (*B*) are considered to deposit into the LB core, with the concentration of these deposited aggregates represented by [*D*_*FR*_] :

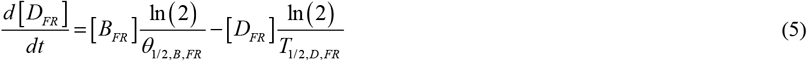

Here, [*D*_*FR*_] represents the concentration of lipid membrane aggregates deposited into the LB core, while *T*_1/ 2, *D, FR*_ is the half-life of these deposited aggregates.

In Eq. (5), the first term on the right-hand side mirrors the fourth term in Eq. (4), but with the opposite sign, indicating the deposition process. The second term accounts for the degradation of lipid fragment aggregates within the LB core, reflecting their possible finite half-life. This term anticipates potential future therapies aimed at LB clearance, possibly through autophagy, as discussed in ref. [41].

Equations (3)–(5) are solved under the following initial conditions:

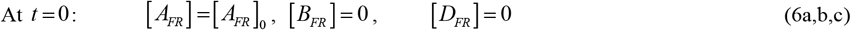

where [*A*_*FR*_]_0_ represents the initial concentration of lipid membrane fragments within the soma at the start of the process.

### 2.2. Analytical solutions for the limiting case of slow deposition of free lipid membrane aggregates into the LB core, *θ*_1/2,*B,FR*_ →∞ (and also *T*_1/ 2, *A, FR*_ →∞, *T*_1/ 2, *B, FR*_ →∞, and *T*_1/ 2, *D, FR*_ →∞)

If the deposition rate of free lipid membrane aggregates into the LB core is slow and the half-lives of monomers, free aggregates, and deposited aggregates are infinitely long, Eqs. (3)-(5) simplify to the following reduced forms:

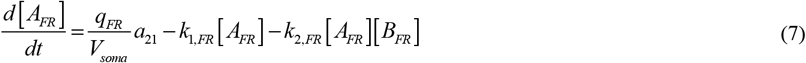

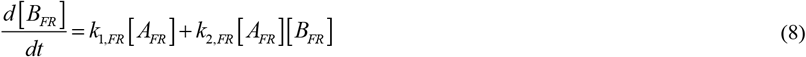

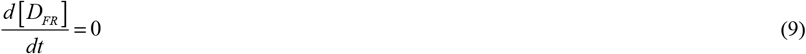

To streamline the analysis, it is assumed that the initial concentrations of lipid membrane fragments are zero – that is,

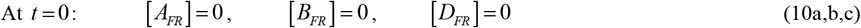

The numerical solution of Eqs. (7)–(10) shows that [*A*_*FR*_] → 0 as *t* →∞. Based on this observation, an approximate solution for Eqs. (7) and (8), valid for large *t*, was derived in ref. [16] as follows:

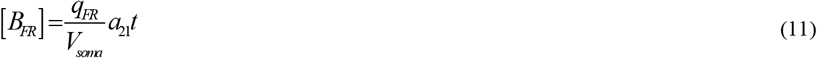

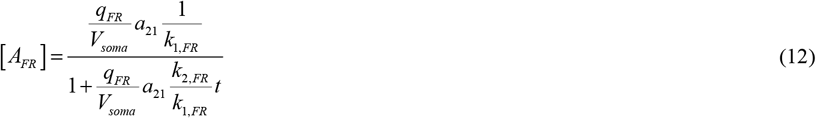

Also, from Eqs. (9) and (10c) it follows that:

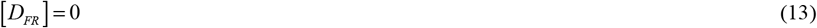

### 2.3. Method for simulating LB core growth

The growth of the LB core (Fig. 1) is determined by calculating the total number of membrane fragments incorporated into the core, *N*_*LB*_, over time *t*. This is done using the following approach, adapted from ref. [42]:

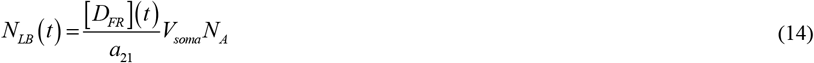

where *N*_*A*_ is Avogadro’s number.

Alternatively, *N*_*LB*_ (*t*) can be determined using the following equation, as outlined in ref. [42]:

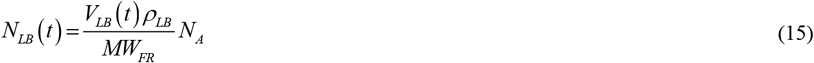

Here *MW*_*FR*_ represents the average molecular weight of a lipid membrane fragment, calculated as the sum of the atomic weights of all atoms within the fragment, *V*_*LB*_ (*t*) is the volume of the LB core at time *t*, and *ρ*_*LB*_ denotes the density of the LB, assumed to be the same for both the core and halo regions.

Equating the right-hand sides of Eqs. (14) and (15) and solving for the LB core volume yields:

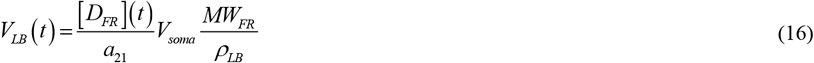

Assuming the LB core maintains a spherical geometry, the volume of the core can be expressed as:

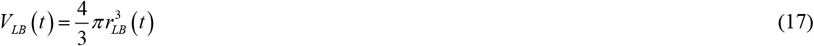

where *r*_*LB*_ represents the radius of the LB core.

Solving Eqs. (16) and (17) for the core radius yields:

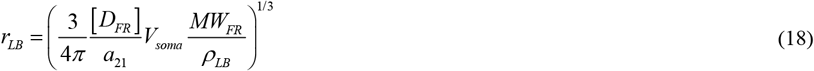

By substituting Eq. (13) into Eq. (18), the following result is obtained:

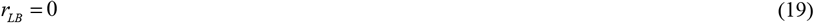

However, if it is assumed that all free membrane fragment aggregates eventually deposit into the core of the LB (noting that this differs from the assumption that *θ*_1/2,*B,FR*_ →∞), the term [*D*_*FR*_] in Eq. (18) can be substituted by [*B*_*FR*_] from Eq. (11), resulting in:

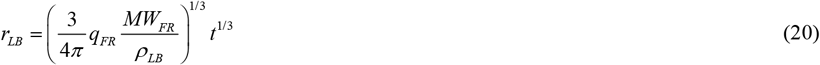

The LB core grows until time *t*_*LB, f*_, at which point *r*_*LB*_ reaches *r*_*LB, f*_. Solving Eq. (20) for *q*_*FR*_ under these conditions gives:

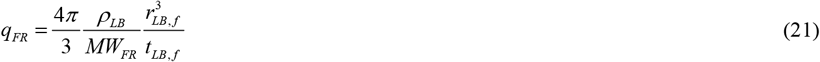

Eq. (21) is essential for estimating model parameters, as it allows for the calculation of *q*_*FR*_ when *r*_*LB, f*_ and *t*_*LB,f*_ are known. This equation estimates *q*_*FR*_ to be 1.57×10^−28^ mol s^−1^, as listed in Table S2.

### 2.4. Equations for simulating the aggregation of *α*-syn monomers in the LB halo

A constant production rate of α-syn monomers in the soma, *q*_*AS*_, is assumed. The F-W model [26], detailed in Eqs. (3)-(5), is employed to simulate the formation of α-syn fibrils in the soma. Stating the conservation of α-syn monomers in the soma gives:

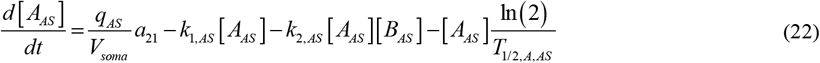

where [*A*_*AS*_] represents the concentration of α-syn monomers in the soma, [*B*_*AS*_] denotes the concentration of free α-syn aggregates in the soma, *k*_1, *AS*_ is the rate constant for the nucleation of free α-syn aggregates, *k*_2, *AS*_ is the rate constant for the autocatalytic growth of these aggregates, and *T*_1/ 2, *A, AS*_ is the half-life of α-syn monomers. Hereafter the words free α-syn aggregates and α-syn oligomers will be used as synonyms.

In Eq. (22), the first term on the right-hand side accounts for the production rate of α-syn monomers. The second and third terms describe the transformation of α-syn monomers into free aggregates via nucleation and autocatalytic growth. The fourth term reflects the decay of α-syn monomers due to their finite half-life.

Applying the conservation to free α-syn aggregates yields:

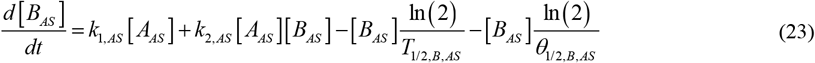

where *θ*_1/ 2, *B, AS*_ represents the half-deposition time of free α-syn aggregates into fibrils.

In Eq. (23), the first two terms represent the increase in free α-syn aggregates due to nucleation and autocatalytic growth, while the third term models their degradation based on their half-life. The fourth term in Eq. (23) represents the reduction in free α-syn aggregates as they are deposited into fibrils. Eqs. (22) and (23) are analogous to Eqs. (3) and (4), respectively.

The process of α-syn fibril formation in the LB halo from free α-syn aggregates is modeled similarly to colloidal suspension coagulation [40]. In this approach, free α-syn aggregates (*B*) are assumed to deposit into fibrils that constitute the LB halo, with their concentration denoted by [*D*_*AS*_]. This results in an equation similar to Eq. (5):

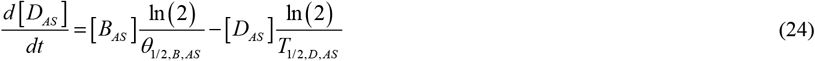

Here [*D*_*AS*_] denotes the concentration of α-syn aggregates deposited into fibrils in the LB halo, and *T*_1/ 2, *D, AS*_ represents the half-life of these deposited aggregates. In Eq. (24), the first term on the right-hand side corresponds to the fourth term in Eq. (23) but with the opposite sign, reflecting the deposition of free aggregates. The second term models the degradation of α-syn aggregates within the fibrils if their half-life is finite. This term also accounts for potential therapeutic interventions, such as autophagy-based therapies [41], which may aim to clear LBs.

Eqs. (22)–(24) are solved under the following initial conditions:

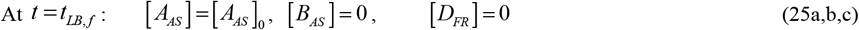

Here [*A*_*AS*_]_0_ represents the initial concentration of α-syn monomers in the soma.

### 2.5. Analytical solutions for the limiting case of slow deposition of free *α*-syn aggregates into fibrils, *θ*_1/2,*B, AS*_ →∞ (and also *T*_1/ 2, *A, AS*_ →∞, *T*_1/ 2, *B, AS*_ →∞, and *T*_1/ 2, *D, AS*_ →∞)

In the case where the deposition rate of free α-syn aggregates into fibrils is slow and the half-lives of α-syn monomers, free aggregates, and deposited aggregates are assumed to be infinitely large, the governing equations (22)–(25) simplify significantly. Under these assumptions, the behavior of the system is reduced to a more manageable form, yielding the following simplified expressions:

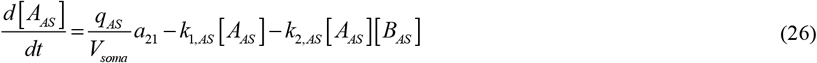

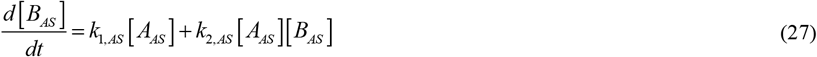

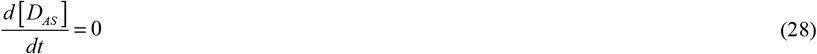

To simplify the analysis, the initial concentration of α-syn monomers in the soma is set to zero, leading to the following initial conditions:

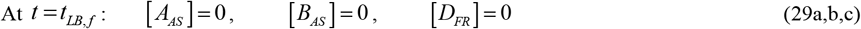

Numerical analysis of Eqs. (26)–(29) indicates that as time increases, [*A*_*AS*_] → 0 as *t* →∞. Based on this observation, an approximate solution for Eqs. (26) and (27), applicable for large values of *t*, was derived in ref. [16] as follows:

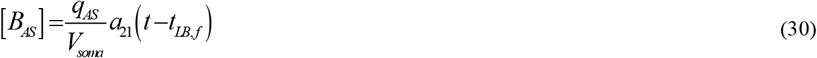

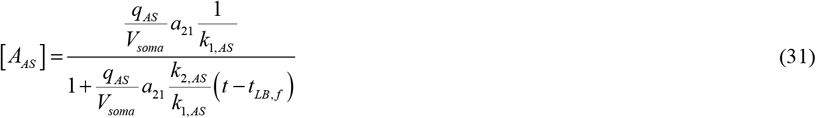

Also, from Eqs. (28) and (29c) it follows that:

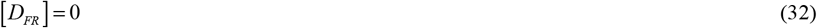

### 2.6. Method for simulating the growth of the LB halo

To model the growth of the halo surrounding the LB core (Fig. 1), the total number of α-syn monomers incorporated into the halo at time *t*, denoted as *N*_*AS*_, is calculated using the following approach [42]:

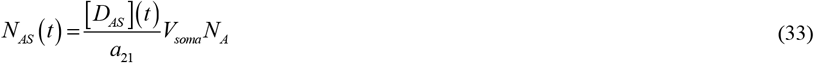

Alternatively, *N*_*AS*_ (*t*) can be determined using the equation provided in ref. [42]:

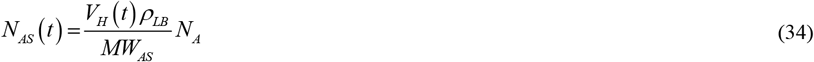

where *MW*_*AS*_ denotes the molecular weight of an α-syn monomer and *V*_*H*_ represents the volume of the LB halo at time *t*.

By equating the right-hand sides of Eqs. (33) and (34) and solving for the LB volume, the following result is obtained:

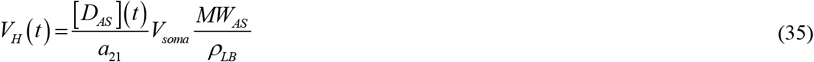

Assuming the LB halo is shaped like a spherical shell,

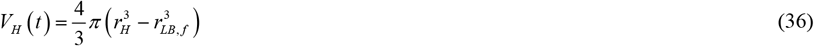

Solving Eqs. (35) and (36) for the halo radius *r*_*H*_ yields:

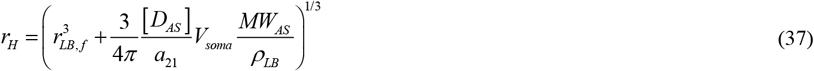

Assuming that free α-syn aggregates eventually deposit into the LB halo (which differs from the assumption that *θ*_1/2,*B, AS*_ →∞), [*D*_*AS*_] in Eq. (37) can be replaced by [*B*_*AS*_], as defined by Eq. (30). Doing so gives:

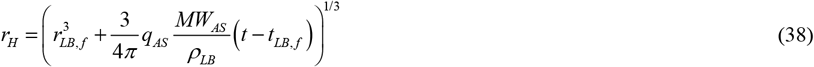

The LB halo will keep growing until it reaches its maximum radius *r*_*H, f*_, at which point *t* equals *t*_*LB, f*_ ± *t*_*H, f*_. To find *q*_*AS*_ Eq. (38) was solved for these values of *r* and *t*, resulting in:

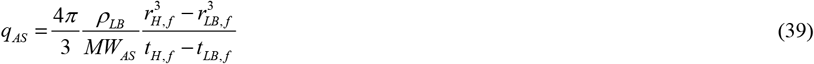

Eq. (39) is crucial for estimating model parameters because it allows for the calculation of *q*_*AS*_ when the values of *r*_*LB, f*_, *r*_*H, f*_, and *t*_*H, f*_ are known. Using Eq. (39), *q*_*AS*_ was estimated to be 1.47×10^−21^ mol s^−1^, as reported in Table S2.

### 2.7. Criterion for assessing accumulated toxicity of *α*-syn oligomers

This paper proposes a parameter to characterize the accumulated toxicity of α-syn oligomers:

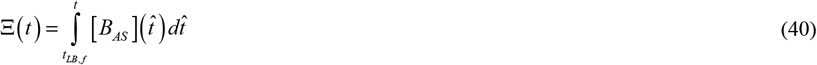

Note that for *t* < *t*_*LB, f*_, toxicity is zero. The dimensionless form of Eq. (40) is given by Eq. (S1) in Supplemental Materials.

Using the approximate Eq. (30) in Eq. (40), it follows that for *t* > *t*_*LB, f*_,

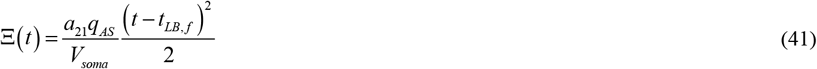

The quadratic relationship between toxicity and time implies that early in the growth of the halo, the accumulated toxicity increases gradually, but over time, the rate of increase becomes significantly faster. Eq. (41) is only valid as *θ*_1/2,*B,AS*_ →∞, *T*_1/ 2, *A, AS*_ →∞, *T*_1/ 2, *B, AS*_ →∞, and *T*_1/ 2, *D, AS*_ →∞.

In the case where *θ*_1/2,*B,AS*_ →0, Ξ also approaches zero. The toxicity of α-syn oligomers (considered equivalent to free aggregates in the model) tends to zero because the aggregates immediately deposit into fibrils, bypassing the stage in which they remain in the cytosol as free aggregates.

### 2.8. Sensitivity of accumulated toxicity of *α*-syn oligomers to model parameters

This study aimed to assess how the accumulated toxicity of α-syn oligomers varies with different model parameters. This analysis involved calculating the local sensitivity coefficients, which are the first-order partial derivatives of accumulated toxicity with respect to parameters such as the half-deposition time of free α-syn aggregates into fibrils, *θ*_1/2,*B,AS*_. The methodology used for this analysis was based on the approaches outlined in refs. [43-46]. Specifically, the sensitivity coefficient of Ξ to a parameter *θ*_1/ 2, *B, AS*_ was determined by:

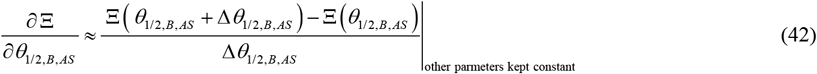

where Δ*θ*_1/ 2, *B, AS*_ = 10^−3^ *θ*_1/ 2, *B, AS*_ represents the step size. To ensure the sensitivity coefficients were independent of the chosen step size, calculations were performed using various step sizes.

The non-dimensional relative sensitivity coefficients were calculated using the method outlined in refs. [44,47], for example,

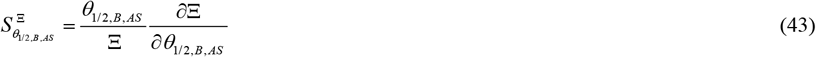

By utilizing Eq. (41), the result can be expressed as follows:

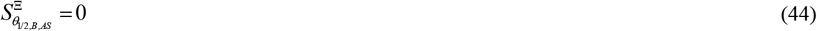

Additionally, utilizing Eq. (41), the sensitivity of Ξ to various parameters is calculated as follows:

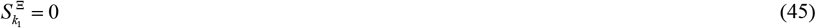

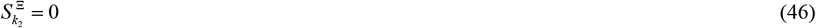

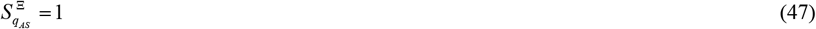

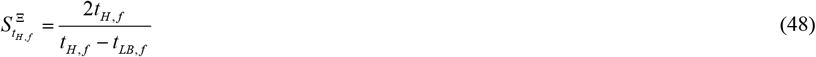

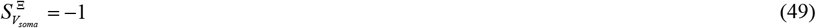

## 3. Results

Details of the numerical solution techniques can be found in section S2 of the Supplemental Materials. In all figures, except for Fig. 5, the following half-lives are used for membrane fragments, free membrane aggregates, α-syn monomers, and α-syn aggregates: *T*_1/ 2, *A, F R*_ = 2. 7 0 × 10 ^5^ s, *T*_1/ 2, *B, FR*_ = 1.35 × 10^6^ s,*T*_1/ 2, *A, AS*_ = 5.76 × 10^4^ s, and *T*_1/ 2, *B, AS*_ = 2.88 × 10^5^ s. The half-lives of deposited membrane aggregates in the LB core and deposited α-syn aggregates in the LB halo are assumed to be infinitely long. For numerical implementation, this is simulated by setting *T*_1/ 2, *D, FR*_ and *T*_1/ 2, *D, FR*_ to 10^20^ s. All other parameter values are as listed in Table 2 unless otherwise specified in the figure or its caption.

First, the numerical solutions for various values of *k*_1_ were examined. The simulations assume that *k*_1,*FR*_ = *k*_1,*AS*_. The molar concentrations of lipid membrane fragments and α-syn monomers, [*A*_*FR*_] and [*A*_*AS*_], respectively, quickly reach equilibrium, remaining constant over time (Fig. S1a). A similar pattern is observed for the molar concentrations of free lipid membrane aggregates in the LB core and free α-syn aggregates, [*B*_*FR*_] and [*B*_*AS*_], respectively, which also reach their equilibrium values and become time-independent (Fig. S1b). The sudden changes in concentrations at *t* = *t*_*LB, f*_ reflect the difference between the molar concentrations of membrane fragments and α-syn.

The molar concentration of lipid membrane aggregates deposited in the LB core, [*D*_*FR*_], remains low during the core’s growth (Fig. S2a), while the concentration of α-syn aggregates deposited in the halo’s fibrils, [*D*_*AS*_], shows a linear increase over time (Fig. S2a). The radii of both the growing LB core and halo follow an approximate cube-root dependency on time (Fig. S2b, the cube root dependency was first established in ref. [16]). The noticeable increase in the LB core radius, *r*_*LB*_, over time, seen in Fig. S2b, appears to contradict the nearly zero concentration of lipid membrane aggregates in the core, [*D*_*FR*_], shown in Fig. S2a. However, a detailed plot of the lipid membrane aggregates deposited into the LB core (Fig. S3a) reveals that these concentrations increase with time, though their values are small, so their increase is not visible in Fig. S2a. This is due to the large average molecular weight of membrane fragments (Table S2). The linear increase in [*D*_*FR*_] over time (Fig. S3a) leads to a corresponding linear increase in the LB core volume, resulting in the core radius growing proportionally to the cube root of time (Fig. 3b). A smaller value of *k*_1_ leads to a smaller LB radius (Figs. S2b and S3b).

Next, the solutions to the problem for various values of *k*_2_ are examined. The simulations assume that *k*_2,*FR*_ = *k*_2,*AS*_. The molar concentrations of lipid membrane fragments, [*A*_*FR*_], remain small (Fig. S4a), while the molar concentrations of α-syn monomers in the halo, [*A*_*AS*_], reach different equilibrium values, depending on *k*_2, *FR*_ and *k*_2, *AS*_ (Fig. S4a). Similarly, the molar concentrations of free lipid membrane aggregates and free α-syn aggregates, [*B*_*FR*_] and [*B*_*AS*_], respectively, follow the same pattern (Fig. S4b).

Interestingly, as *k*_2, *FR*_ and *k*_2, *AS*_ increase, the concentration of α-syn monomers in the halo decreases, whereas the concentration of free α-syn aggregates in the halo increases.

The molar concentration of lipid membrane aggregates deposited into the LB core, [*D*_*FR*_], remains small during the core’s growth (Fig. S5a), while the concentration of α-syn aggregates deposited in the halo’s fibrils, [*D*_*AS*_], shows a linear increase over time (Fig. S5a). The radii of the growing LB core and halo, *r*_*LB*_ and *r*_*H*_, respectively, follow a cube-root dependency with time (Fig. S5b), with smaller values of *k*_2_ corresponding to a smaller LB radius (Figs. S5b).

The effects of the production rates of lipid membrane fragments and α-syn monomers are similar to the effects of *k*_2_. The concentrations of free α-syn aggregates, [*B*_*AS*_], reach different equilibrium levels depending on the value of *q*_*AS*_ (Fig. S6b). Meanwhile, the concentration of α-syn aggregates deposited in the halo’s fibrils, [*D*_*AS*_], increases linearly over time, with faster growth observed for larger *q*_*AS*_ values (Fig. S7a). The radii of both the growing LB core and the halo, *r*_*LB*_ and *r*_*H*_, expand approximately according to a cube root dependence with time, with faster growth for higher *q*_*FR*_ and *q*_*AS*_ values (Fig. S7b).

The accumulated toxicity of α-syn oligomers, Ξ, calculated using Eq. (40), increases linearly over time and is independent of *k*_1_ (Fig. 2a). This linear increase occurs because the concentration of α-syn oligomers, [*B*_*AS*_], remains constant over time (Fig. S1b) and 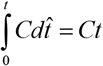 if *C* is constant. While the radius of the LB halo, *r*_*LB*_, also increases over time, it does so nonlinearly (Fig. 2b). The dependence of the halo radius on *k*_1_ in Fig. 2b may seem unexpected since [*D*_*AS*_] in Fig. S2 is independent of *k*_1_, but the LB core radius is influenced by *k*_1_ (Fig. S2b), causing the halo to begin expanding from different initial radii that depend on the value of *k*_1_.

**Fig. 2.**
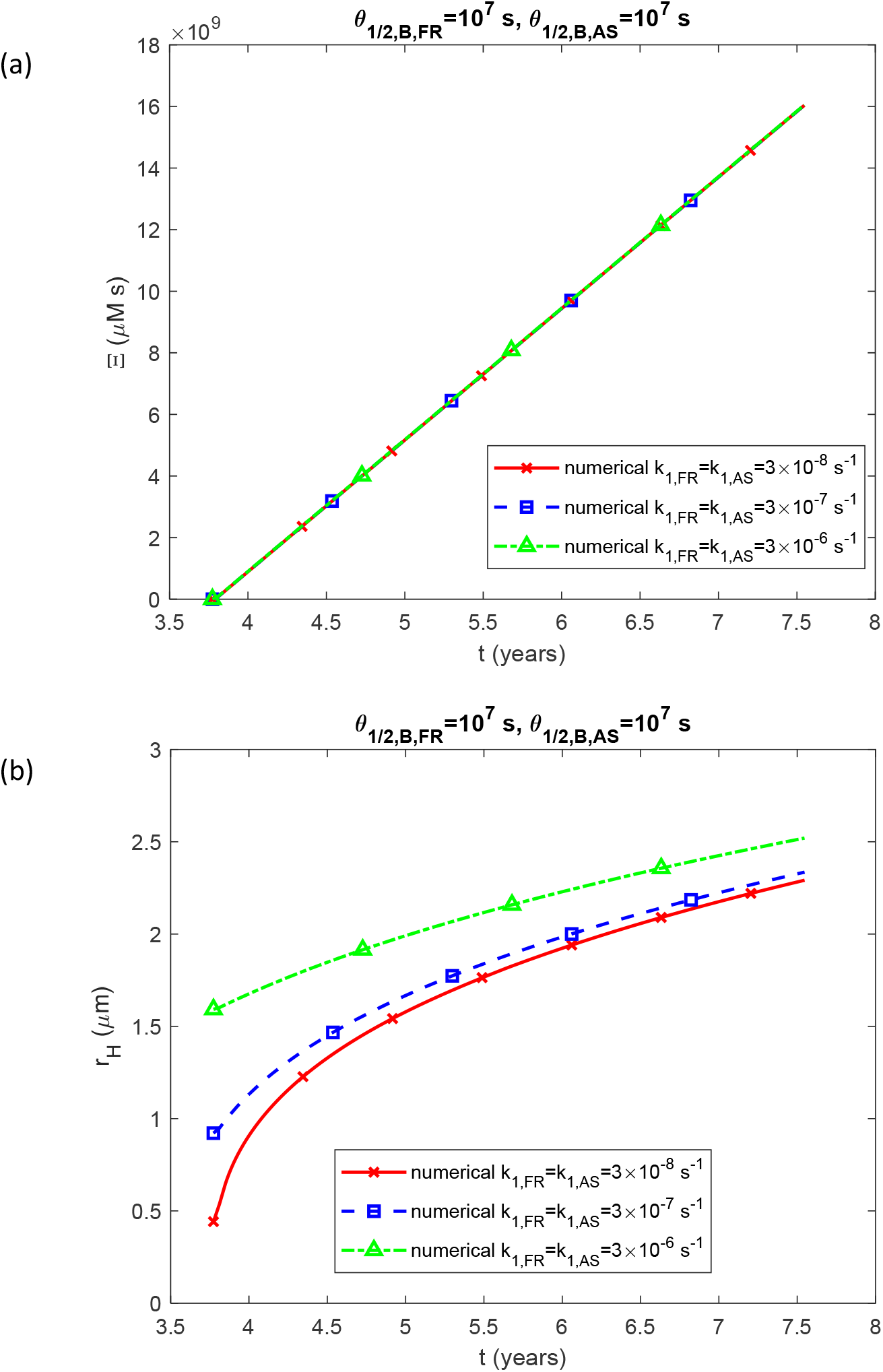
*(a) A*ccumulated toxicity of α-syn oligomers, Ξ, vs time. (b) The radius of the growing LB halo, *r*_*H*_, vs time. The results are presented for various values of *k*_1, *FR*_ and *k*_1, *AS*_. (*k*_2,*FR*_ = *k*_2, *AS*_ = 2×10^−6^ μM^−1^ s^−1^, *q*_*FR*_ =1.57×10^−28^ mol s^−1^, *q*_*AS*_ = 1.47 × 10^−21^ mol s^−1^.)

The accumulated toxicity rises as *k*_2_ increases (Fig. 3a), with the halo radius showing a similar growth trend (Fig. 3b). Likewise, the accumulated toxicity also increases when the production rates of membrane fragments and α-syn monomers, *q*_*FR*_ and *q*_*AS*_, are higher (Fig. 4a). The halo radius also increases when *q*_*FR*_ and *q*_*AS*_ increase (Fig. 4b). The scenario shown in Fig. 4 was recalculated using half-lives of monomers and free aggregates that are an order of magnitude shorter, corresponding to a tenfold increase in their degradation rates (*T*_1/ 2, *A, F R*_ = 2. 7 0 × 10 ^4^ s, *T*_1/ 2, *B, FR*_ = 1.35 × 10^5^ s, *T*_1/ 2, *A, AS*_ = 5.76 × 10^3^ s, *T*_1/ 2, *B, AS*_ = 2.88 × 10^4^ s). The results indicate approximately 15 times lower toxicity (Fig. 5a) and a roughly 2.5 times smaller LB halo radius (Fig. 5b).

**Fig. 3.**
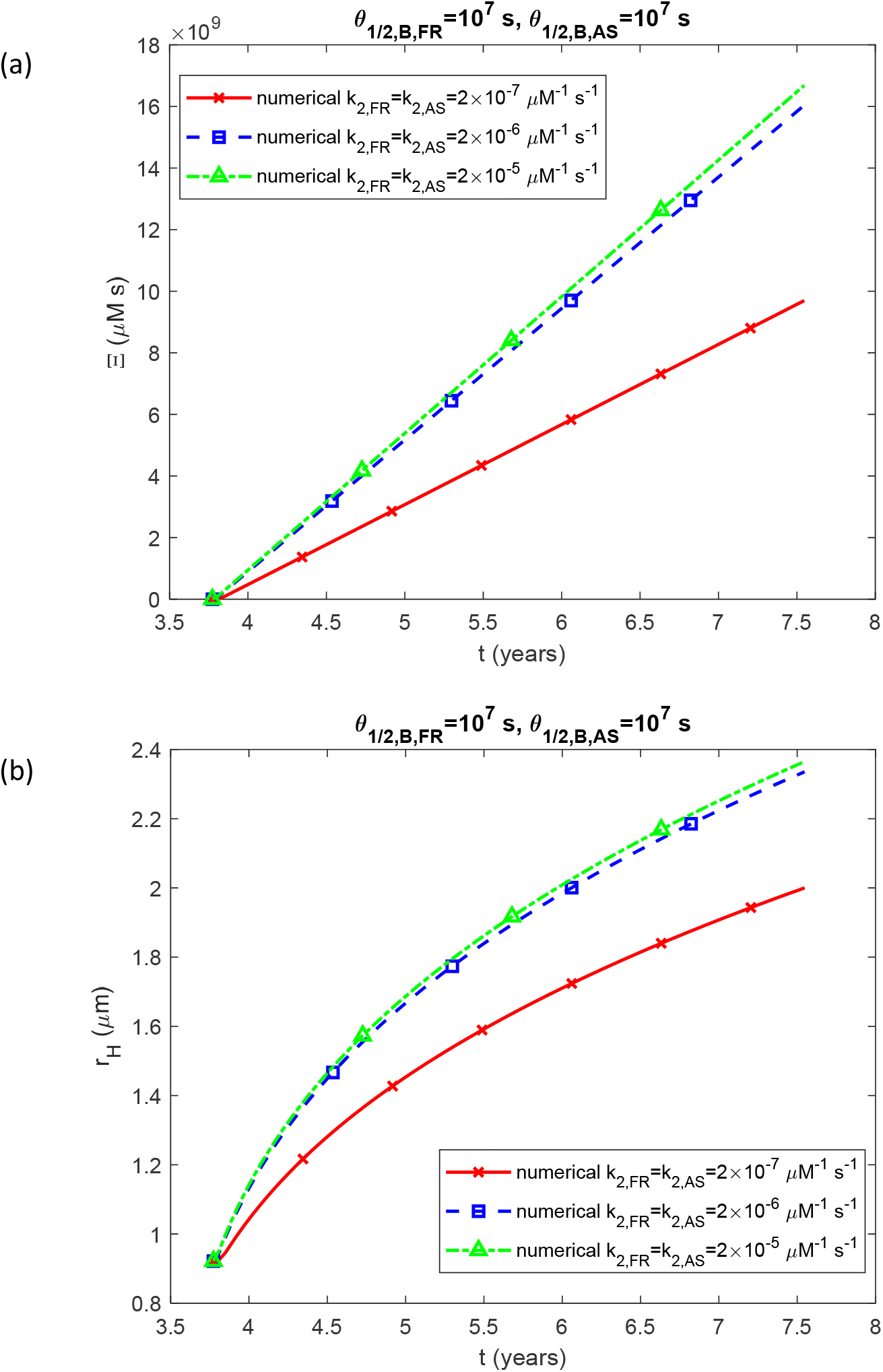
(a) Accumulated toxicity of α-syn oligomers, Ξ, vs time. (b) Radius of the growing LB halo, *r*_*H*_, vs time. Results are displayed for different values of *k*_2, *FR*_ and *k*_2, *AS*_ (*k*_1,*FR*_ = *k*_1, *AS*_ = 3×10^−7^ s^−1^, *q*_*FR*_ =1.57×10^−28^ mol s^−1^, *q*_*AS*_ = 1.47 × 10^−21^ mol s^−1^.)

**Fig. 4.**
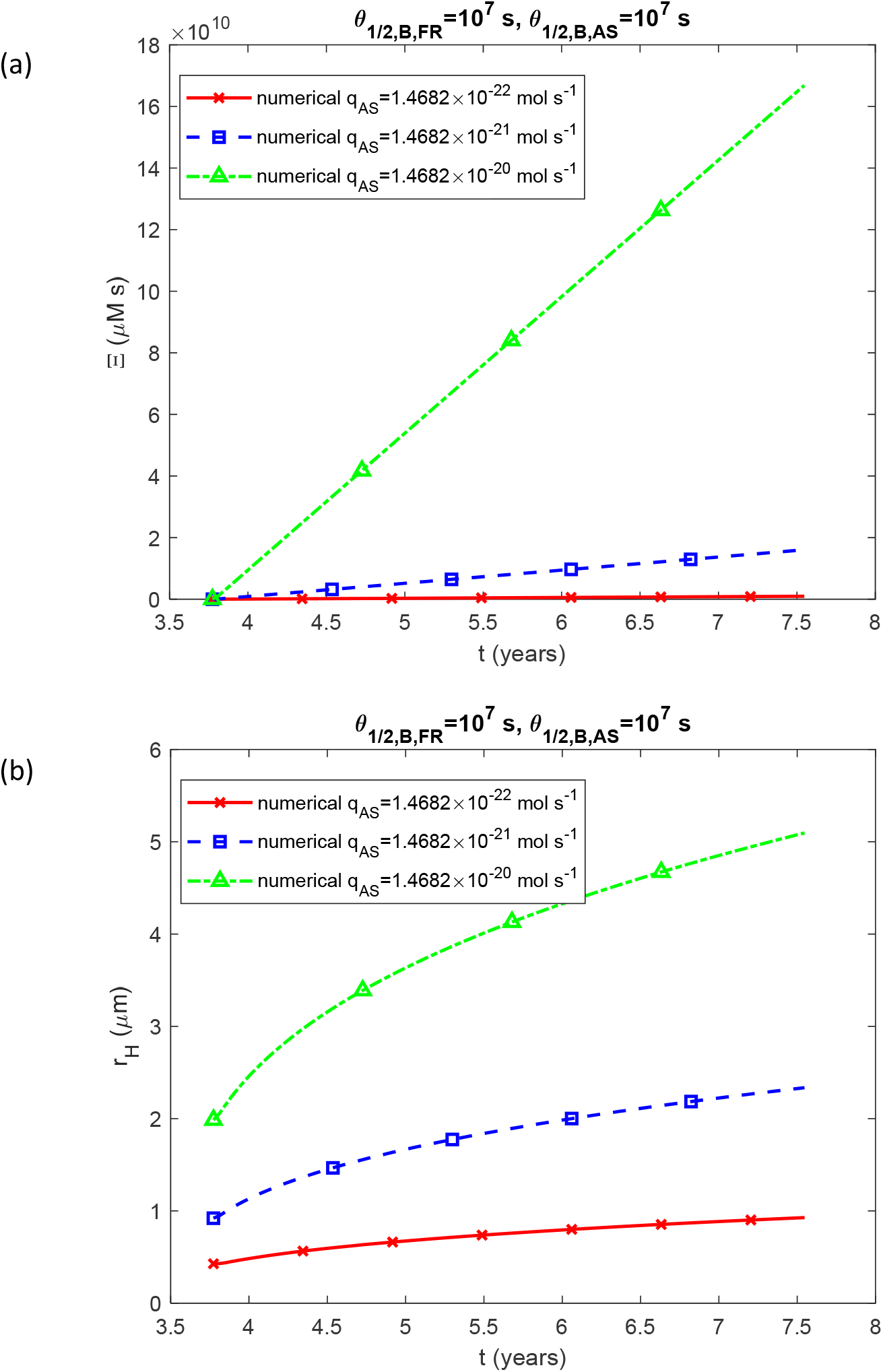
(a) Accumulated toxicity of α-syn oligomers, Ξ, vs time. (b) The radius of the expanding LB halo vs time. Results are presented for various values of *q*_*FR*_ and *q*_*AS*_. The following corresponding values were used for *q*_*FR*_ : *q*_*FR*_ =1.57×10^−29^ mol s^−1^ was used for *q*_*AS*_ =1.47×10^−22^ mol s^−1^, *q*_*FR*_ =1.57×10^−28^ mol s^−1^ was used for *q*_*AS*_ =1.47×10^−21^ mol s^−1^, and *q*_*FR*_ =1.57×10^−27^ mol s^−1^ was used for *q*_*AS*_ =1.47×10^−20^ mol s^−1^. (*k*_1,*FR*_ = *k*_1, *AS*_ = 3×10^−7^ s^−1^, *k*_2,*FR*_ = *k*_2, *AS*_ = 2×10^−6^ μM^−1^ s^−1^.)

**Fig. 5.**
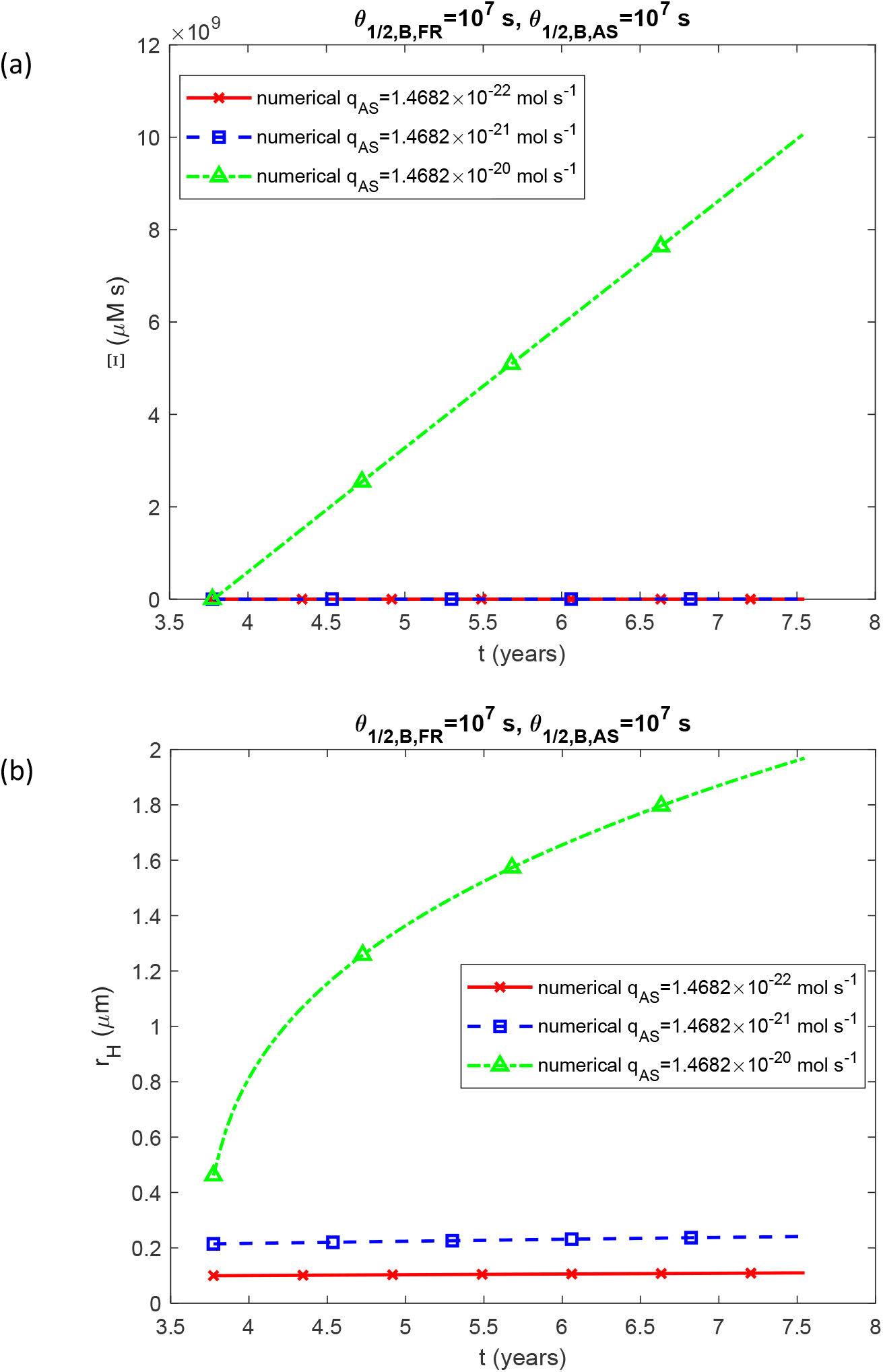
Similar to Fig. 4, but with monomers and free aggregates having half-lives reduced by an order of magnitude: *T*_1/ 2, *A, FR*_ = 2.70 × 10^5^ s, *T*_1/ 2, *B, FR*_ = 1.35 × 10^6^ s, *T*_1/ 2, *A, AS*_ = 5.76 × 10^4^ s, *T*_1/ 2, *B, AS*_ = 2.88 × 10^5^ s.

The role of LBs in PD remains unclear, with researchers debating whether they merely sequester toxic α-syn oligomers or are themselves harmful to neurons [12-14,30,48,49]. If LBs simply act as storage for cytotoxic α-syn species, one would expect their numbers to increase as the disease progresses.

Surprisingly, this is not the case. As PD advances, the number of neurons containing LBs in the substantia nigra pars compacta does not rise; instead, neurons harboring LBs tend to die [15]. This observation points to a more active role for LBs in contributing to neurodegeneration. The increase in accumulated toxicity of α-syn aggregates may be the true cause of neuron death, with the size of LBs merely showing a correlation with this toxicity.

Fig. 6a-c suggests a correlation between accumulated toxicity, Ξ, and the radius of the LB halo, *r*_*H*_, such that as *r*_*H*_ increases, Ξ also increases. Assuming that neurons die when the halo radius reaches its maximum value, the critical level of accumulated toxicity can be roughly estimated from Fig. 6a-c as 10^10^ µM·s. However, this is a preliminary estimate that requires validation through future experimental studies. It is important to note that the curves representing Ξ(*r*_*H*_) for small values of *q*_*AS*_ in Fig. 6c terminate at lower *r*_*H*_ values. This is because the halo radius does not grow significantly for small *q*_*AS*_ (Figs. 4b and S7b).

**Fig. 6.**
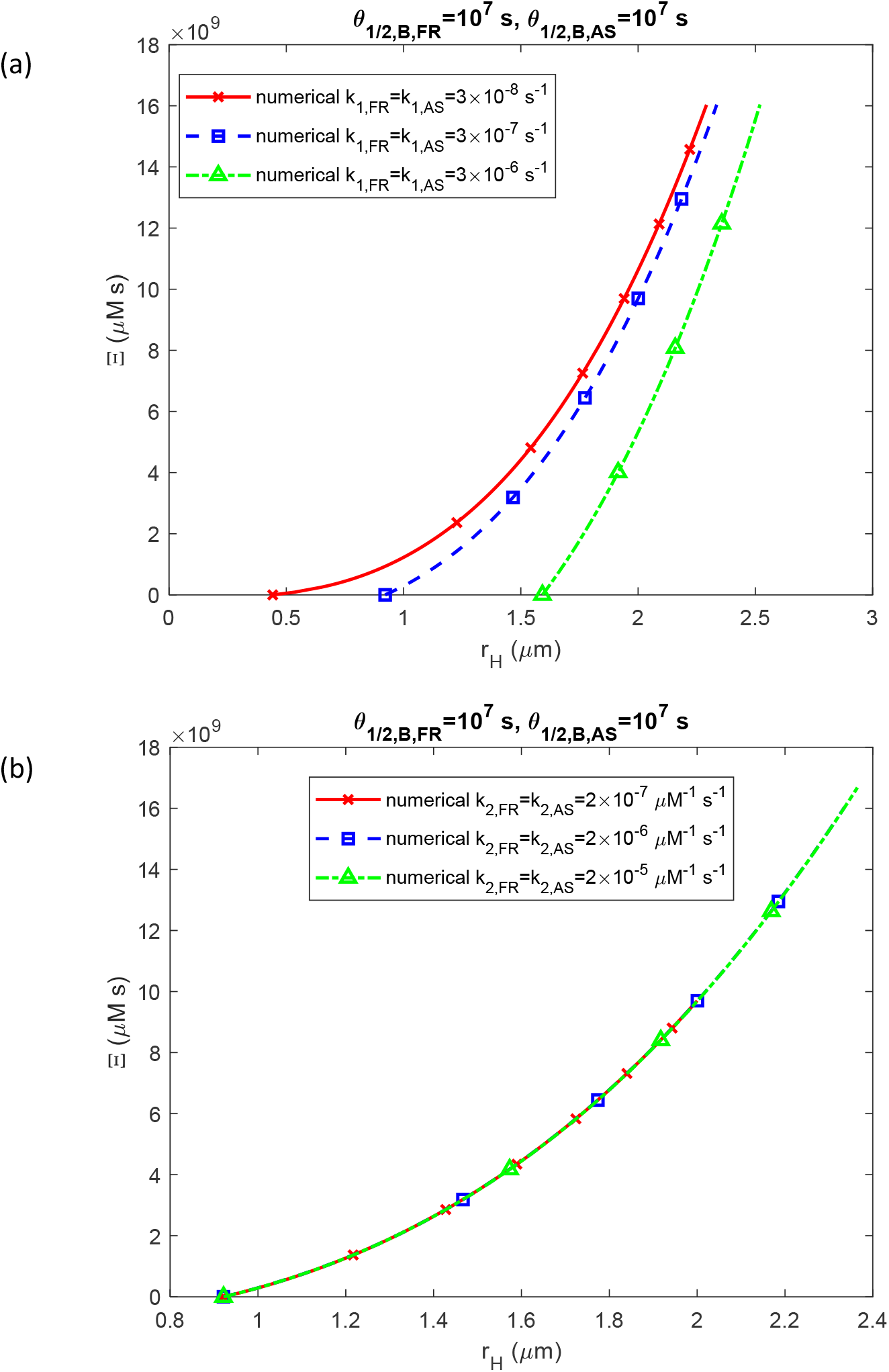
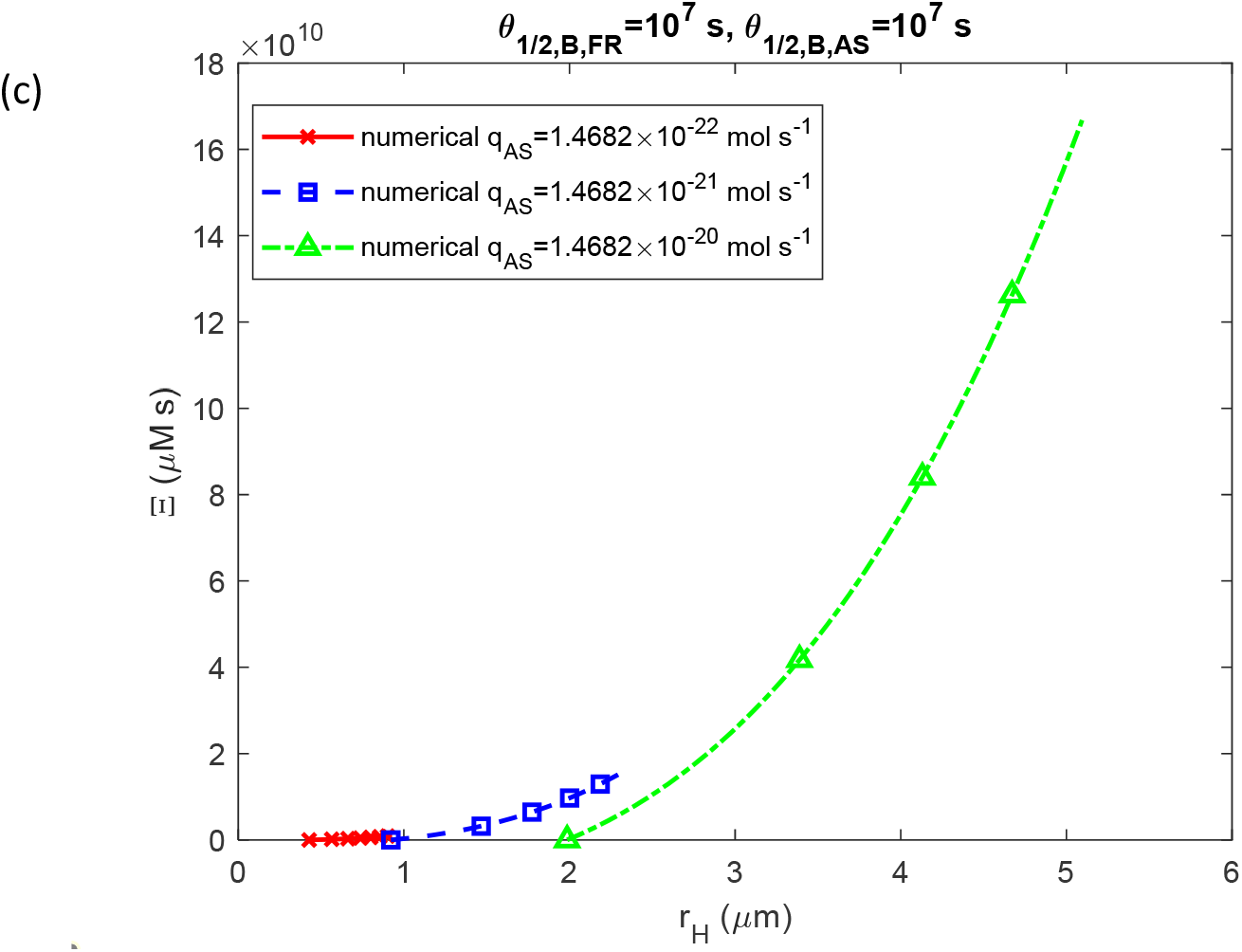
The accumulated toxicity of α-syn oligomers, Ξ, vs the radius of a growing LB halo, *r*_*H*_, is shown for three different conditions: (a) varying nucleation rate constants, *k*_1, *FR*_ and *k*_1, *AS*_, (b) varying autocatalytic rate constants, *k*_2, *FR*_ and *k*_2, *AS*_, and (c) different production rates of membrane fragments and α-syn monomers, *q*_*FR*_ and *q*_*AS*_. The following corresponding values were used for *q*_*FR*_ : *q*_*FR*_ =1.57×10^−29^ mol s^−1^ was used for *q*_*AS*_ =1.47×10^−22^ mol s^−1^, *q*_*FR*_ =1.57×10^−28^ mol s^−1^ was used for *q*_*AS*_ =1.47×10^−21^ mol s^−1^, and *q*_*FR*_ =1.57×10^−27^ mol s^−1^ was used for *q*_*AS*_ = 1.47×10^−20^ mol s^−1^. (*k*_1,*FR*_ = *k*_1, *AS*_ = 3×10^−7^ s^−1^, *k*_2,*FR*_ = *k*_2, *AS*_ = 2×10^−6^ μM^−1^ s^−1^.)

When *θ*_1/2,*B,AS*_ is small, α-syn oligomers rapidly deposit into fibrils, resulting in zero accumulated toxicity (Fig. 7a-c). This supports the hypothesis that LBs may have a neuroprotective function [7]. The highest level of accumulated toxicity occurs as *θ*_1/2,*B,AS*_ →∞ because, in this case, free α-syn aggregates remain in the cytosol indefinitely, continuing to catalyze their own production from α-syn monomers. The model suggests that deposition into LBs effectively removes α-syn oligomers from the cytosol. Notably, some lines in Fig. 7a-c cross the critical toxicity threshold, 10^10^ µM s, when *θ*_1/2,*B,AS*_ increases. This is an indication that these neurons will die.

**Fig. 7.**
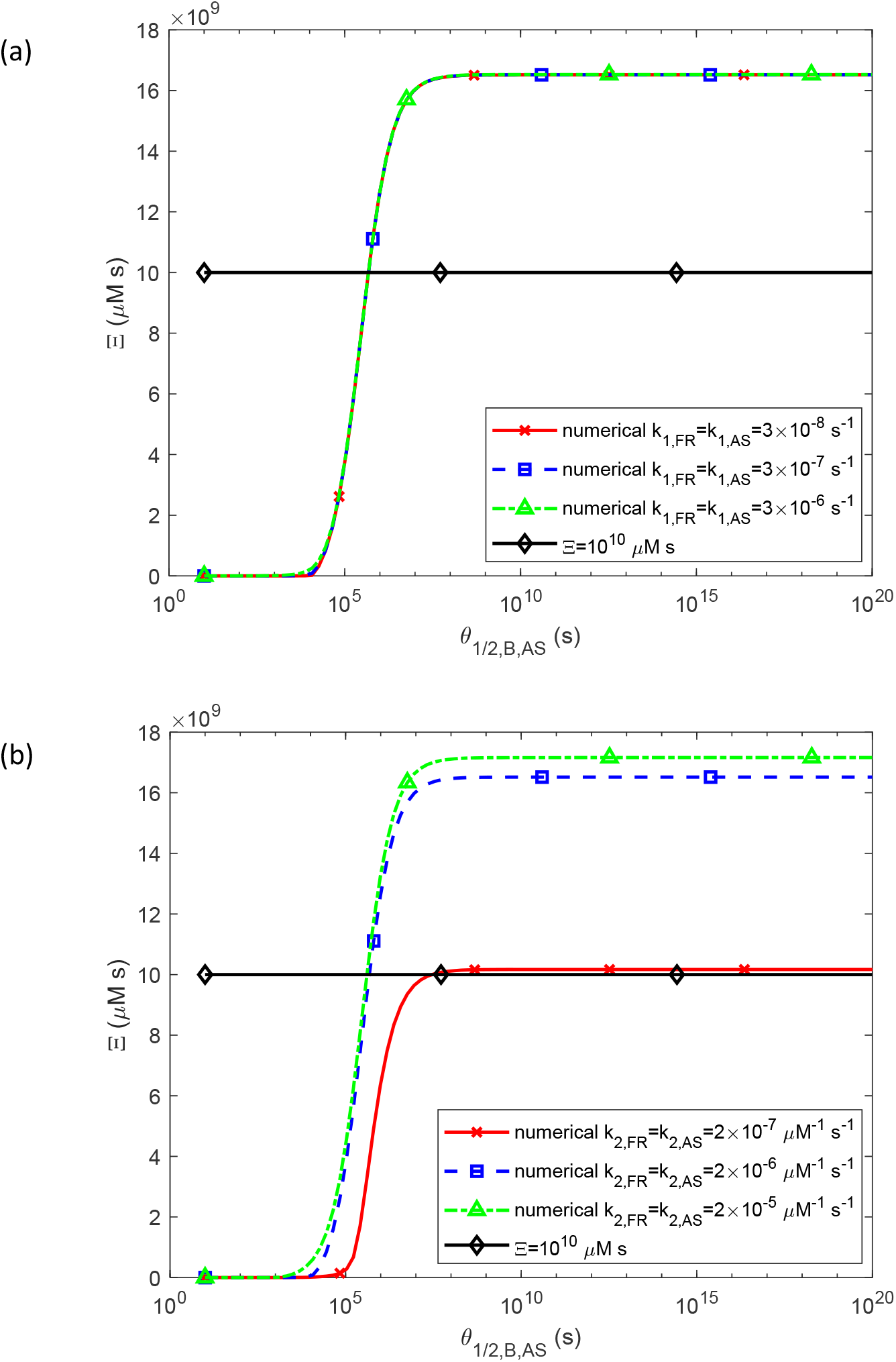
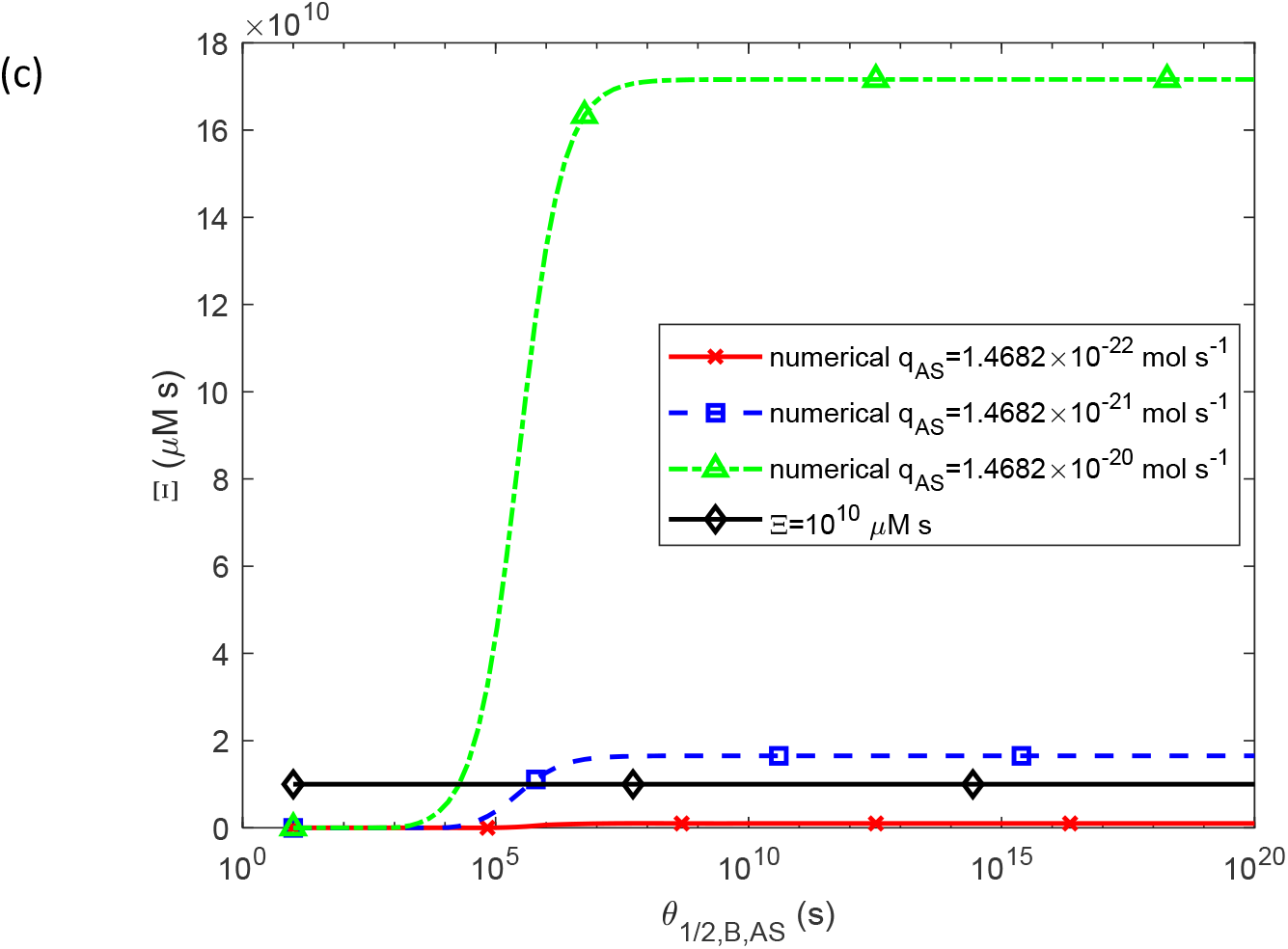
The accumulated toxicity of α-syn oligomers, Ξ, vs the half-deposition time of free α-syn aggregates into fibrils, *θ*_1/2,*B,AS*_, for three different values of (a) nucleation rate constants, *k*_1, *FR*_ and *k*_1, *AS*_ (*k*_2,*FR*_ = *k*_2, *AS*_ = 2×10^−6^ μM^−1^ s^−1^, *q*_*AS*_ =1.57×10^−28^ mol s^−1^, *q*_*FR*_ = 1.47 × 10^−21^ mol s^−1^), (b) autocatalytic rate constants, *k*_2, *FR*_ and *k*_2, *AS*_ (*k*_1,*FR*_ = *k*_1, *AS*_ = 3×10^−7^ s^−1^, *q*_*FR*_ =1.57×10^−28^ mol s^−1^, *q*_*AS*_ = 1.47 × 10^−21^ mol s^−1^), (c) production rates of membrane fragments and α-syn monomers, *q*_*FR*_ and *q*_*AS*_. The following corresponding values were used for *q*_*FR*_ : *q*_*AS*_ =1.57×10^−29^ mol s^−1^ was used for *q*_*FR*_ =1.47×10^−22^ mol s^−1^, *q*_*FR*_ =1.57×10^−28^ mol s^−1^ was used for *q*_*AS*_ =1.47×10^−21^ mol s^−1^, and *q*_*FR*_ =1.57×10^−27^ mol s^−1^ was used for *q*_*AS*_ =1.47×10^−20^ mol s^−1^. (*k*_1,*FR*_ = *k*_1, *AS*_ = 3×10^−7^ s^−1^, *k*_2,*FR*_ = *k*_2, *AS*_ = 2×10^−6^ μM^−1^ s^−1^.) An assumption that *θ*_1/2,*B,FR*_ =*θ*_1/2,*B,AS*_ was used in the computer code.

The curves showing the dependence of Ξ on *θ*_1/2,*B,AS*_ are unaffected by the kinetic constants related to nucleation processes, *k*_1, *FR*_ and *k*_1, *AS*_ (Fig. 7a). However, Ξ increases as the kinetic constants of the autocatalytic process, *k*_2, *FR*_ and *k*_2, *AS*_, increase (Fig. 7b), indicating that these autocatalytic constants have a greater impact on accumulated toxicity than the nucleation constants, *k*_1, *FR*_ and *k*_1, *AS*_. Additionally, accumulated toxicity increases rapidly with increased production rates of membrane fragments and α-syn monomers, *q*_*FR*_ and *q*_*AS*_ (Fig. 7c).

The sensitivity of accumulated toxicity, Ξ, to the half-deposition time of free α-syn aggregates into fibrils, *θ*_1/2,*B,AS*_, reveals sharp peaks in the transition region (Fig. 8a-c). This transition occurs from a zero value of Ξ (when free α-syn aggregates rapidly deposit into LBs) to the maximum value of Ξ at large *θ*_1/2,*B,AS*_ (when free α-syn aggregates remain in the cytosol and do not deposit into LBs) (Fig. 7a-c). The location of the peak of 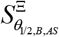 is unaffected by the values of *k*_1,*FR*_ and *k*_1, *AS*_, though the amplitude of the peak increases as *k*_1, *AS*_ decreases (Fig. 8a). Both the amplitude and location of the peak of 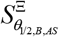 remain independent of *k*_2,*FR*_ and *k*_2, *AS*_ (Fig. 8b). The peak location of sensitivity shifts to smaller *θ*_1/2,*B,AS*_ values as *q*_*AS*_ increases. Interestingly, for the smallest *q*_*AS*_ value, the peak of 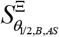 disappears entirely (Fig. 8c).

**Fig. 8.**
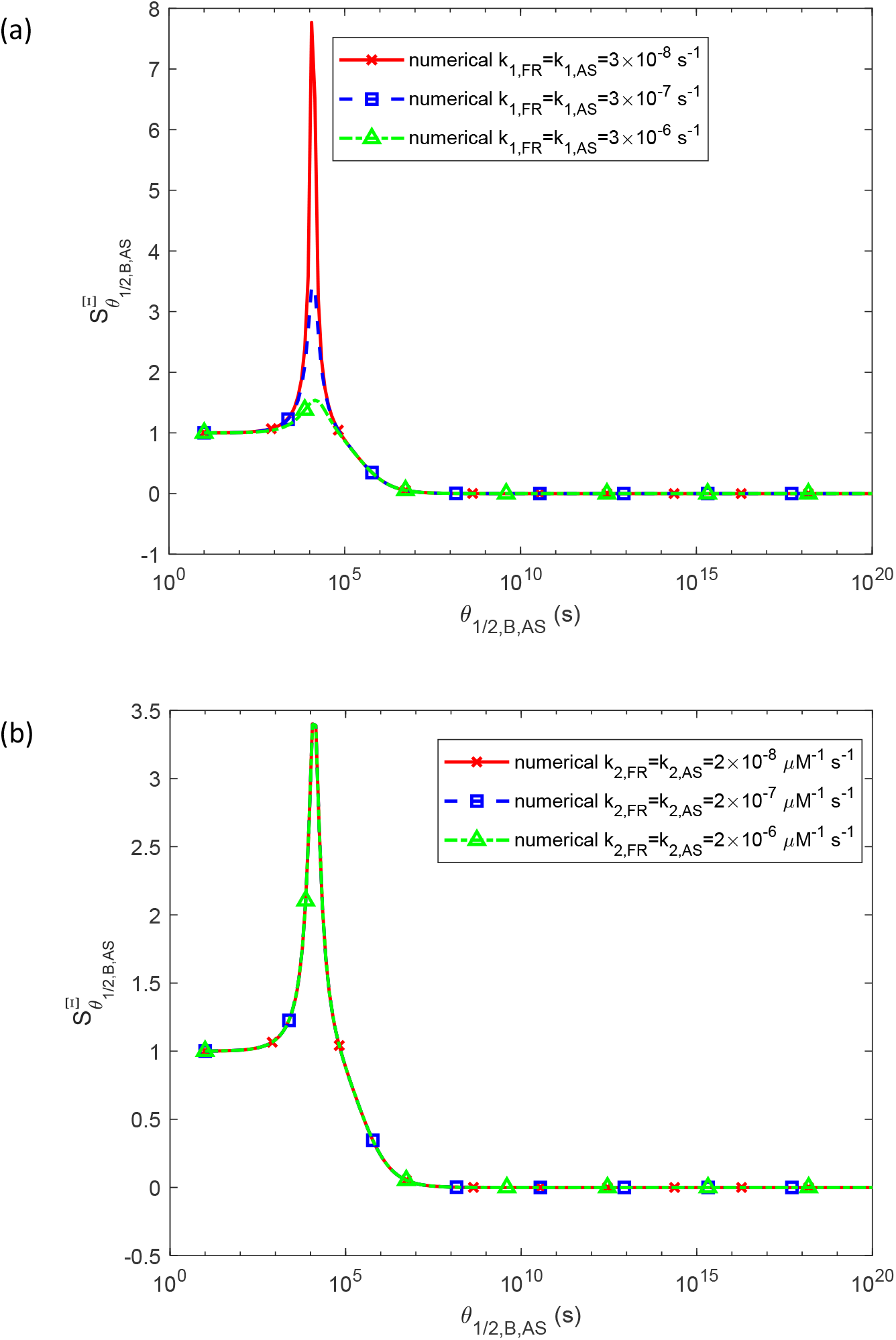
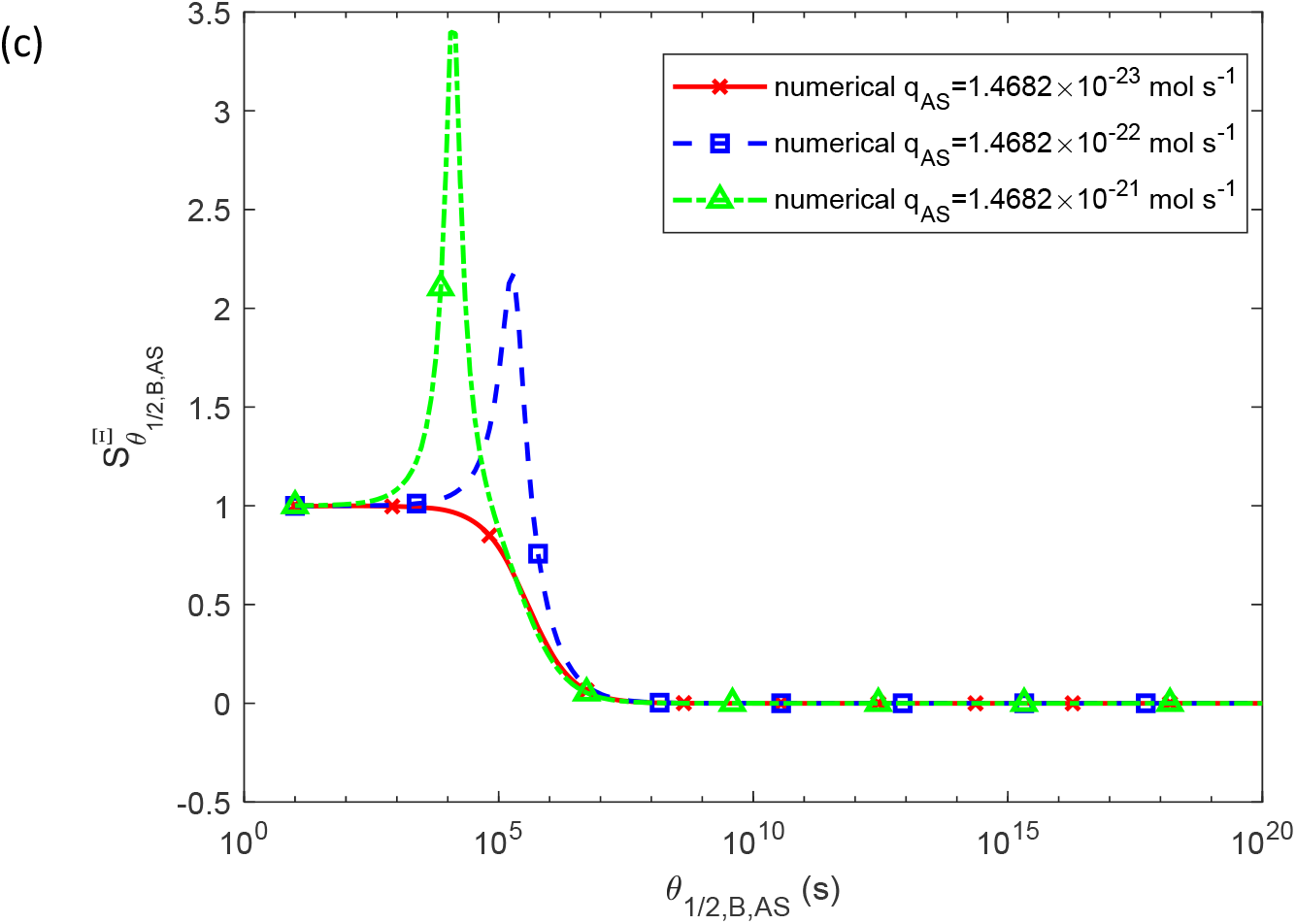
Sensitivity of accumulated toxicity, Ξ, to the half-deposition time of free α-syn aggregates into fibrils, *θ*_1/2,*B,AS*_, is examined for three different values of (a) the rate constants that describe the nucleation of membrane fragments and α-syn aggregates, *k*_1,*FR*_ and *k*_1, *AS*_ (*k*_2,*FR*_ = *k*_2, *AS*_ = 2×10^−6^ μM^−1^ s^−1^, *q*_*FR*_ =1.57×10^−28^ mol s^−1^, *q*_*AS*_ = 1.47 × 10^−21^ mol s^−1^), (b) the rate constants that describe the autocatalytic growth of membrane fragments and α-syn aggregates, *k*_2,*FR*_ and *k*_2, *AS*_ (*k*_1,*FR*_ = *k*_1, *AS*_ = 3×10^−7^ s^−1^, *q*_*FR*_=1.57×10^−28^ mol s^−1^, *q* _*AS*_= 1.47 × 10^−21^ mol s^−1^), (c) the production rate of α-syn monomers, *q*_*AS*_. (*k*_1,*FR*_ = *k*_1, *AS*_= 3×10^−7^ s^−1^, *k*_2,*FR*_= *k*_2, *AS*_= 2×10^−6^ μM^−1^ s^−1^.) An assumption that *θ*_1/2,*B,FR*_=*θ*_1/2,*B,AS*_ was used in the computer code.

Note that the numerically computed curves for 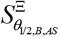 align with the analytical prediction, 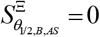, given by Eq. (41), as *θ*_1/2,*B,AS*_ →∞.

Age-related declines in α-syn degradation mechanisms have been proposed as a contributing factor to α-syn aggregation [4]. Malfunctions in these degradation pathways would result in longer half-lives for α-syn monomers and free aggregates, *T*_1/ 2, *A, AS*_ and *T*_1/ 2, *B, AS*_, respectively. An increase in *T*_1/ 2, *A, AS*_ leads to an S-shaped rise in accumulated toxicity, Ξ (Fig. 9). Importantly, an increase in *T*_1/ 2, *A, AS*_ causes the accumulated toxicity to exceed the critical threshold of 10^10^ µM·s, indicating that the affected neurons will undergo cell death. This increase in Ξ is independent of the nucleation rate constants, *k*_1, *FR*_ and *k*_1, *AS*_ (Fig. S9a). Higher values of the rate constants associated with autocatalytic growth, *k*_2, *FR*_ and *k*_2, *AS*_, cause the rise in Ξ to begin at smaller values of *T*_1/ 2, *A, AS*_. However, the asymptotic value of Ξ for large *T*_1/ 2, *A, AS*_ remains unaffected by *k*_2, *FR*_ and *k*_2, *AS*_ (Fig. S9b). Additionally, the asymptotic values of Ξ for large *T*_1/ 2, *A, AS*_ increase with an increase in the production rate of α-syn monomers, *q*_*AS*_ (Fig. S9c).

**Fig. 9.**
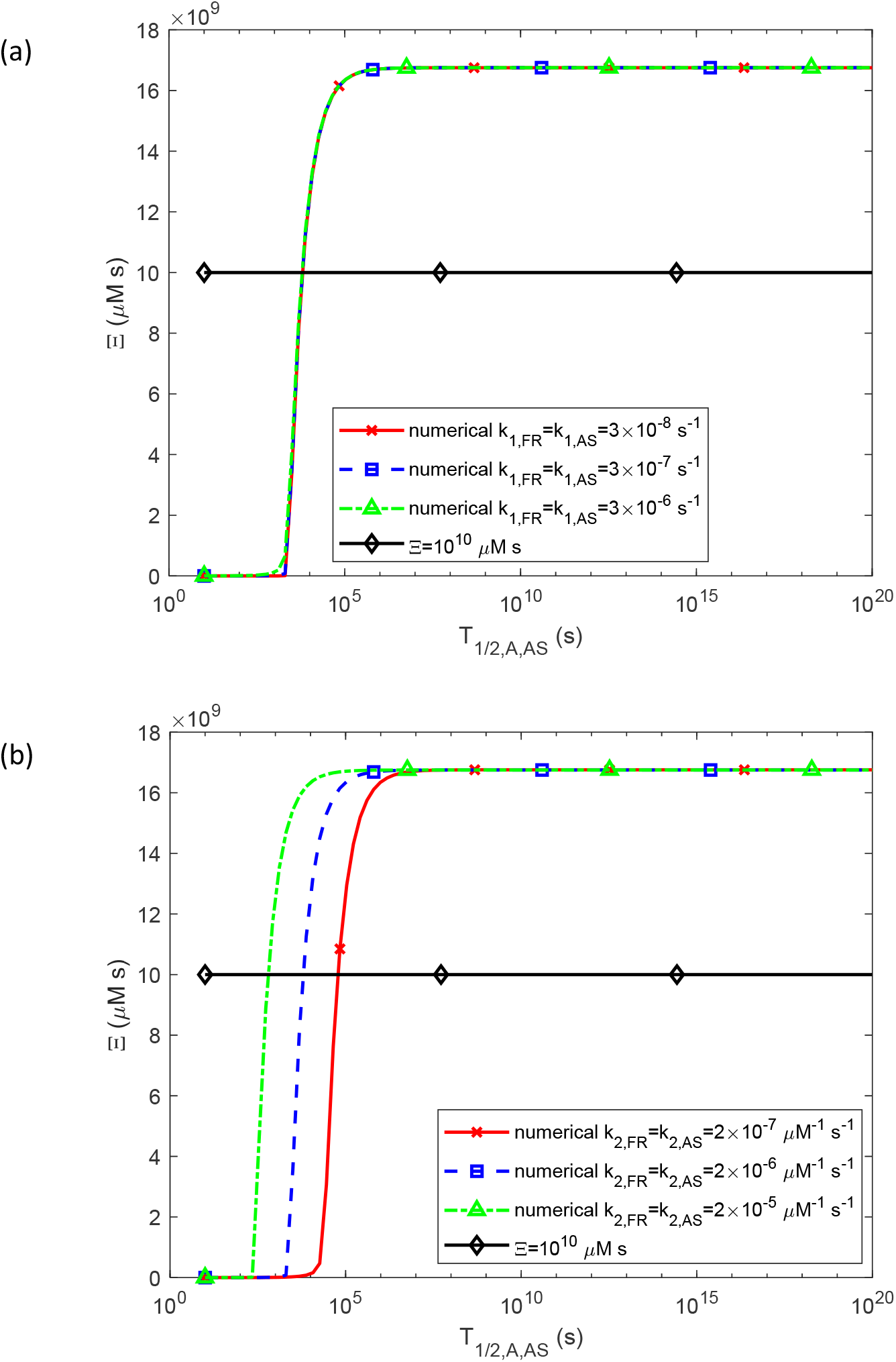
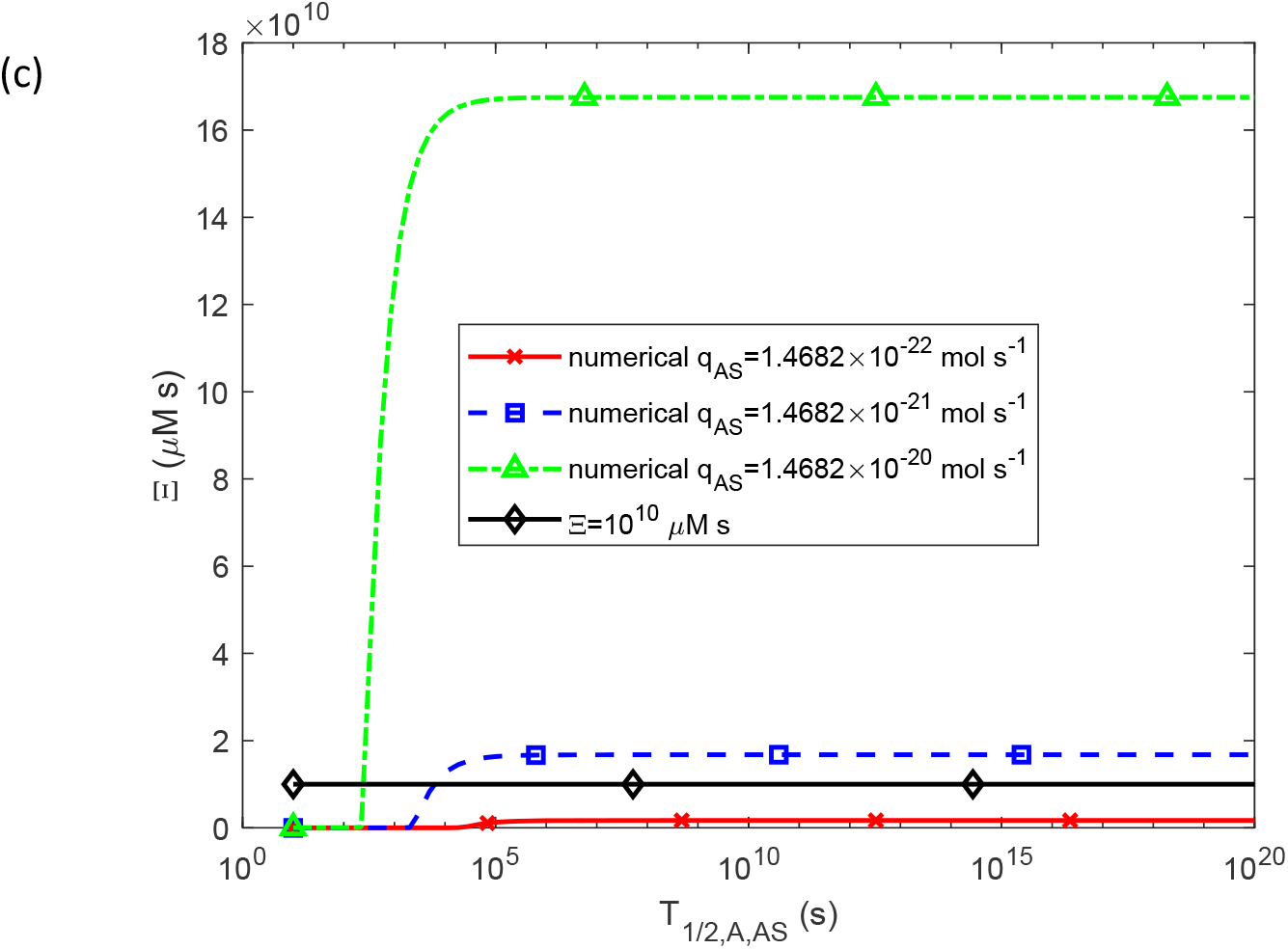
The accumulated toxicity of α-syn oligomers, Ξ, vs the half-life of α-syn monomers, *T*_1/ 2, *A, AS*_, for three different values of (a) nucleation rate constants, *k*_1,*FR*_ and *k*_1, *AS*_ (*k*_2,*FR*_ = *k*_2, *AS*_ = 2×10^−6^ μM^−1^ s^−1^, *q*_*FR*_ =1.57×10^−28^ mol s^−1^, *q* _*AS*_= 1.47 × 10^−21^ mol s^−1^), (b) autocatalytic rate constants, *k*2,*FR* and *k*_2, *AS*_ (*k*_1,*FR*_ = *k*_1, *AS*_ = 3×10^−7^ s^−1^, *q*_*FR*_ =1.57×10^−28^ mol s^−1^, *q*_*AS*_ = 1.47 × 10^−21^ mol s^−1^), (c) production rates of membrane fragments and α-syn monomers, *q*_*FR*_ and *q*_*AS*_. The following corresponding values were used for *q* _*FR*_ : *q*_*FR*_ =1.57×10^−29^ mol s^−1^ was used for *q* _*AS*_ =1.47×10^−22^ mol s^−1^, *q* _*FR*_ =1.57×10^−28^ mol s^−1^ was used for *q*_*AS*_=1.47×10^−21^ mol s^−1^, and *q* _*FR*_=1.57×10^−27^ mol s^−1^ was used for *q* _*AS*_=1.47×10^−20^ mol s^−1^. (*k*_1,*FR*_ = *k*_1, *AS*_ = 3×10^−7^ s^−1^, *k*_2,*FR*_ = *k*_2, *AS*_ = 2×10^−6^ μM^−1^ s^−1^.) *T*_1/ 2, *B, AS*_, *T*_1/ 2, *A, FR*_, and *T*_1/ 2, *B, FR*_ were kept at their values given in Table S2.

The accumulated toxicity, Ξ, also shows an S-shaped increase with the increase in *T*_1/ 2, *B, AS*_ (Fig. 10). The increase in *T*_1/ 2, *B, AS*_ results in the accumulated toxicity exceeding the critical threshold of 10^10^ µM·s. This increase is independent of the values of *k*_1, *FR*_ and *k*_1, *AS*_ (Fig. S10a). There is a minimal dependence of Ξ on the growth of *k*_2, *FR*_ and *k*_2, *AS*_ (Fig. S10b), with smaller values of *k*_2, *FR*_ and *k*_2, *AS*_ resulting in a slightly lower asymptotic value of Ξ as *T*_1/ 2, *B, AS*_ → ∞. A larger value of *q*_*AS*_ as leads to a higher asymptotic value *T*_1/ 2, *B, AS*_ → ∞ (Fig. 10c).

**Fig. 10.**
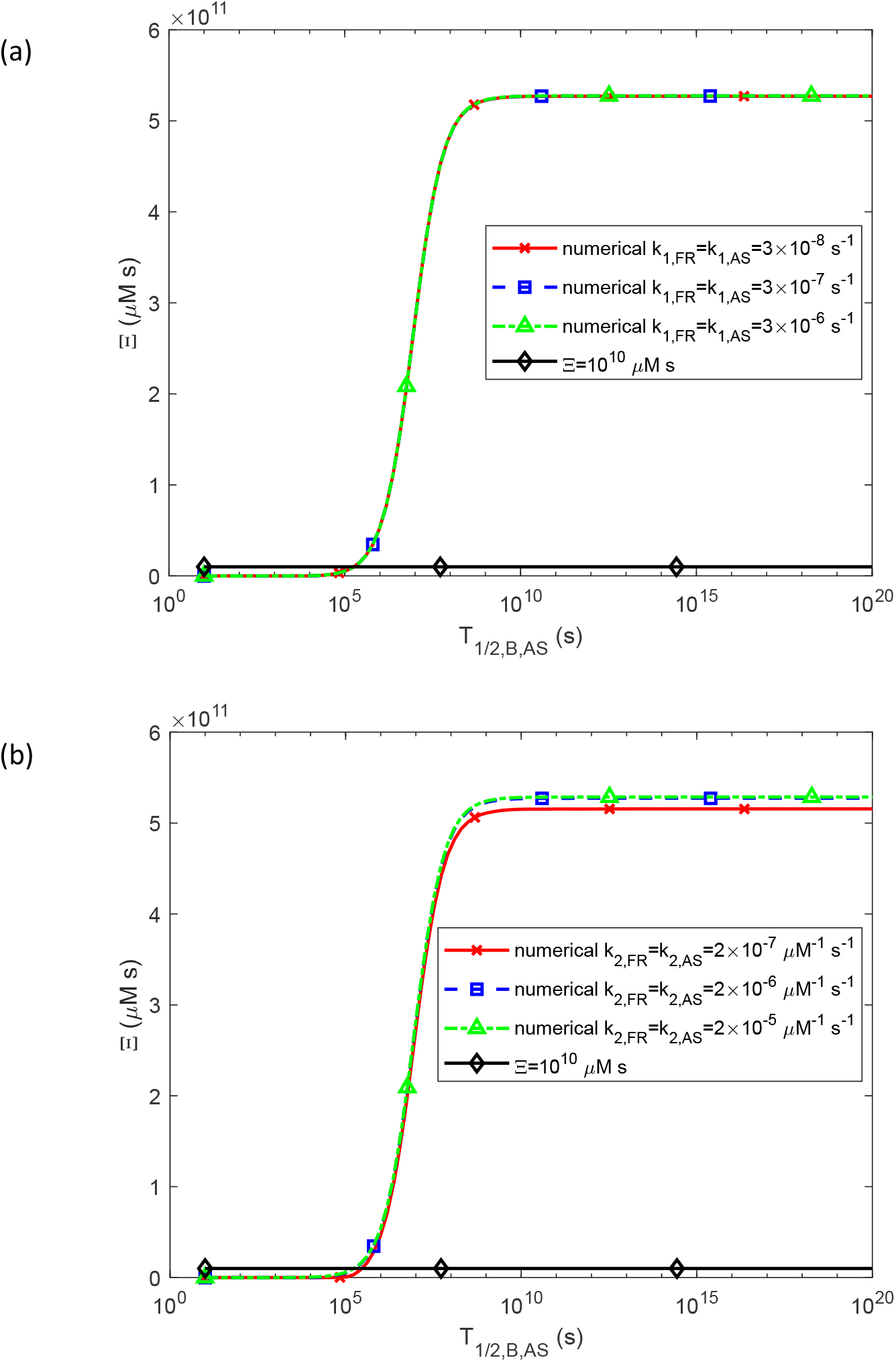
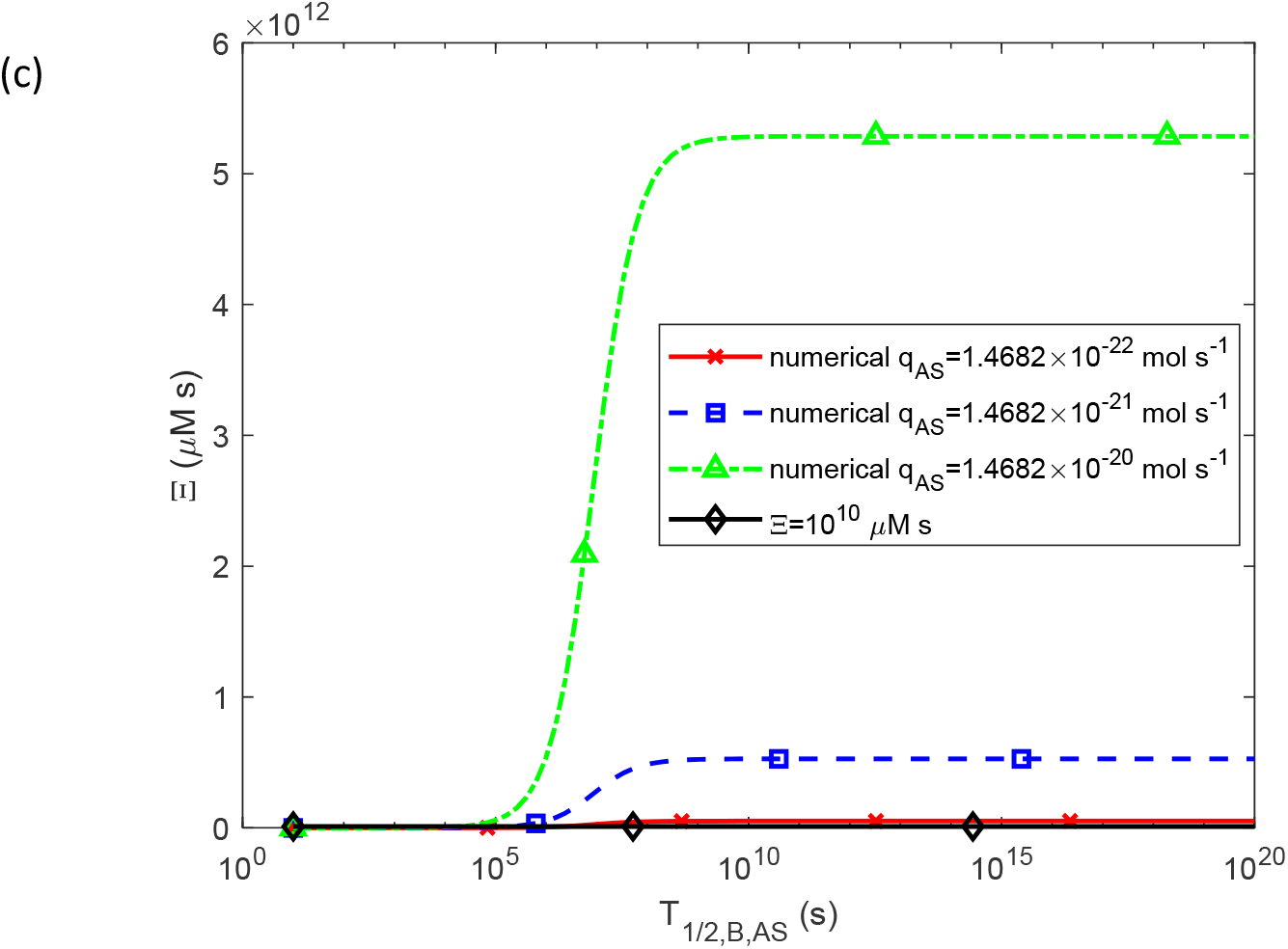
The accumulated toxicity of α-syn oligomers, Ξ, vs the half-life of free α-syn aggregates, *T*_1/ 2, *B, AS*_, for three different values of (a) nucleation rate constants, *k*_1, *FR*_ and *k*_1, *AS*_ (*k*_2,*FR*_ = *k*_2, *AS*_ = 2×10^−6^ μM^−1^ s^−1^, *q*_*FR*_ =1.57×10^−28^ mol s^−1^, *q*_*AS*_ = 1.47 × 10^−21^ mol s^−1^), (b) autocatalytic rate constants, *k*_2, *FR*_ and *k*_2, *AS*_ (*k*_1,*FR*_ = *k*_1, *AS*_ = 3×10^−7^ s^−1^, *q*_*FR*_ =1.57×10^−28^ mol s^−1^, *q*_*AS*_ = 1.47 × 10^−21^ mol s^−1^), (c) production rates of membrane fragments and α-syn monomers, *q*_*FR*_ and *q*_*AS*_. The following corresponding values were used for *q*_*FR*_ : *q*_*FR*_ =1.57×10^−29^ mol s^−1^ was used for *q*_*AS*_ =1.47×10^−22^ mol s^−1^, *q*_*FR*_ =1.57×10^−28^ mol s^−1^ was used for *q*_*AS*_ =1.47×10^−21^ mol s^−1^, and *q*_*FR*_ =1.57×10^−27^ mol s^−1^ was used for *q*_*AS*_ =1.47×10^−20^ mol s^−1^. (*k*_1,*FR*_ = *k*_1, *AS*_ = 3×10^−7^ s^−1^, *k*_2,*FR*_ = *k*_2, *AS*_ = 2×10^−6^ μM^−1^ s^−1^.) *T*_1/ 2, *A, AS*_, *T*_1/ 2, *A, FR*_, and *T*_1/ 2, *B, FR*_ were kept at their values given in Table S2.

The sensitivity of accumulated toxicity, Ξ, to the half-life of α-syn monomers, *T*_1/ 2, *A, AS*_, displays a peak at the point where Ξ(*T*_1/2, *A, AS*_) in Fig. 9 transitions from zero to a high value. The location of these peaks is independent of *k*_1, *FR*_ and *k*_1, *AS*_, although their amplitudes are influenced by *k*_1, *FR*_ and *k*_1, *AS*_ (Fig. S8). The sensitivity peak of Ξ shifts to smaller values of *T*_1/ 2, *A, AS*_ when *k*_2, *FR*_ and *k*_2, *AS*_ increase (Fig. S8b), which aligns with the results shown in Fig. 9b. Additionally, increasing *q* _*AS*_ also shifts the sensitivity peak of 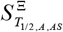 towards smaller values of *T*_1/ 2, *A, AS*_ (Fig. S8c), suggesting that for higher production rates of α-syn monomers, the transition from zero to large values of Ξ occurs at smaller *T*_1/ 2, *A, AS*_, consistent with the results in Fig. 9c. Intriguingly, the curves representing 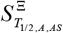 in Fig. S8b for three different values of *k*_2, *AS*_ (differing by an order of magnitude) coincide with the curves in Fig. 8c for three different values of *q*_*AS*_ (also differing by an order of magnitude). This indicates that 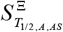 responds in a similar manner to changes in both *k*_2, *AS*_ and *q*_*AS*_.

The position of the peak sensitivity of accumulated toxicity, Ξ, to the half-life of free α-syn aggregates, *T*_1/ 2, *B, AS*_, remains unaffected by changes in *k*_1, *FR*_ and *k*_1, *AS*_. However, the peak amplitude increases as *k*_1, *FR*_ and *k*_1, *AS*_ decrease (Fig. S9a). The position of the peak of 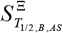 shifts to lower values of *T*_1/ 2, *B, AS*_ when *k*_2, *FR*_ and *k*_2, *AS*_ increase (Fig. S9b). Similarly, an increase in the production rate of α-syn monomers, *q*_*AS*_, also causes the position of the peak of 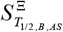 to shift to smaller values of *T*_1/ 2, *B, AS*_ (Fig. S9c). The curves illustrating 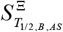 in Fig. S9b for three different values of *k*_2, *AS*_ (differing by an order of magnitude) are identical to those in Fig. 9c for three different values of *q*_*AS*_ (also differing by an order of magnitude). This suggests that 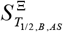 exhibits an identical response to variations in both *k*_2, *AS*_ and *q*_*AS*_.

## 4. Discussion, limitations of the model, and future directions

The numerical investigation indicates that, for physiologically relevant parameter values, the concentration of free α-syn oligomers remains constant over time (Figs. S1b, S4b, and S6b). Since the accumulated toxicity of α-syn oligomers is defined as the integral of their concentration over time, the accumulated toxicity increases linearly with time (Figs. 2a, 3a, and 4a). Assuming neuron death occurs when the radius of the LB halo becomes large, the critical value of accumulated toxicity is estimated to be around 10^10^ µM·s.

The concentration of α-syn aggregates deposited in the LB halo also increases linearly over time (Figs. S2a, S5a, and S7a), leading to a linear increase in the volume of the LB halo. Consequently, the halo radius increases in proportion to the cube root of time (Figs. S2b, S5b, and S7b).

The rate of accumulated toxicity from α-syn oligomers rises with an increase in α-syn monomer production in the soma (Fig. 4a). Shortening the half-lives of monomers and free aggregates leads to a reduction in accumulated toxicity (Fig. 5a). Accordingly, reducing the production rate of α-syn monomers or accelerating their degradation could help slow the accumulation of toxicity.

The results shown in Fig. 6 suggest a strong correlation between accumulated toxicity and the radius of the LB halo. This could explain why neurons die when the LBs they contain grow large (Shults, 2006). The underlying cause may not be the size of the LB itself but rather the high level of accumulated toxicity, which happens to be correlated with the LB size.

Rapid deposition of α-syn oligomers into fibrils results in zero accumulated toxicity (Fig. 7), so LBs may have a neuroprotective role [7]. Conversely, the highest toxicity occurs when oligomers deposit slowly, remaining in the cytosol and catalyzing their own production from monomers. The model suggests that LB formation helps clear α-syn oligomers from the cytosol, reducing their toxic effects.

The accumulated toxicity shows a strong sensitivity to the half-deposition time of free α-syn aggregates into fibrils. A pronounced peak appears in the range where accumulated toxicity shifts from negligible (at short half-deposition times) to high levels (at longer times), compare Figs. 7 and 8. The exact position of this peak varies depending on other parameter values.

The findings demonstrate that increased half-lives of α-syn monomers and aggregates cause an S-shaped increase in accumulated toxicity, which eventually surpasses the critical threshold and leads to neuron death. This aligns with the widely accepted hypothesis in biomedical literature that disruptions in protein degradation pathways may contribute to the progression of PD [21].

α-syn oligomers can form in two ways: either during the self-assembly of α-syn monomers or through the release from already-formed fibrillar species [22]. Future research should expand the model to include the potential release of α-syn oligomers from fibrillar species.

Since misfolded, aggregated species of α-syn are known for their ability to catalyze their own production and propagate to connected neurons (acting as propagons, as noted in ref. [7]), it would be important to develop a model that combines the current framework for α-syn aggregation with one that describes the transport of α-syn aggregates. These aggregates could act as seeds for the formation of new α-syn aggregates in neighboring neurons.

## Abbreviations

α-syn: alpha-synuclein
F-W: Finke-Watzky
LB: Lewy body
PD: Parkinson’s disease

## Acknowledgment

AVK acknowledges the support provided by the National Science Foundation (grant CBET-2042834) and the Alexander von Humboldt Foundation through the Humboldt Research Award.

## Supplemental Materials

### S1. Dimensionless parameter for accumulated toxicity of *α*-syn oligomers

The accumulated toxicity of α-syn oligomers, expressed in Eq. (40), can be reformulated into a dimensionless quantity as follows:

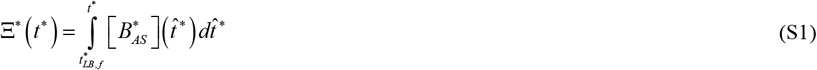

where

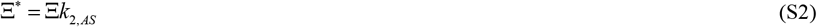

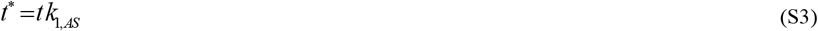

and

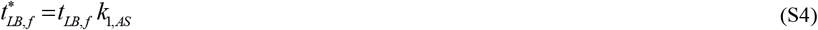

### S2. Numerical Solution

The system of equations, Eqs. (3)-(6) and (24)-(27), was solved using MATLAB’s ODE45 solver (MATLAB R2020b, MathWorks, Natick, MA, USA). To ensure high precision, the relative and absolute tolerance parameters, RelTol and AbsTol, were set to 1e-10.

### S3. Supplementary tables

**Table S1.** Dependent variables in the model.

| Symbol | Definition | Units |
| --- | --- | --- |
| $[A_{FR}]$ | Molar concentration of all fragments within the soma that can contribute to the formation of the LB core—specifically, lipid membrane fragments, damaged organelles, lysosomes, and dysfunctional mitochondria | $\mu\text{M}$ |
| $[B_{FR}]$ | Molar concentration of free lipid membrane aggregates present in the soma | $\mu\text{M}$ |
| $[D_{FR}]$ | Molar concentration of lipid membrane aggregates that have been deposited into the LB core | $\mu\text{M}$ |
| $[A_{AS}]$ | Molar concentration of α-syn monomers present in the soma | $\mu\text{M}$ |
| $[B_{AS}]$ | Molar concentration of unbound α-syn aggregates in the soma | $\mu\text{M}$ |
| $[D_{AS}]$ | Molar concentration of α-syn aggregates that have been deposited into fibrils within the LB halo | $\mu\text{M}$ |
| $r_{LB}$ | Radius of the growing LB core | $\mu\text{m}$ |
| $r_H$ | Radius of the growing LB halo | $\mu\text{m}$ |

**Table S2.**
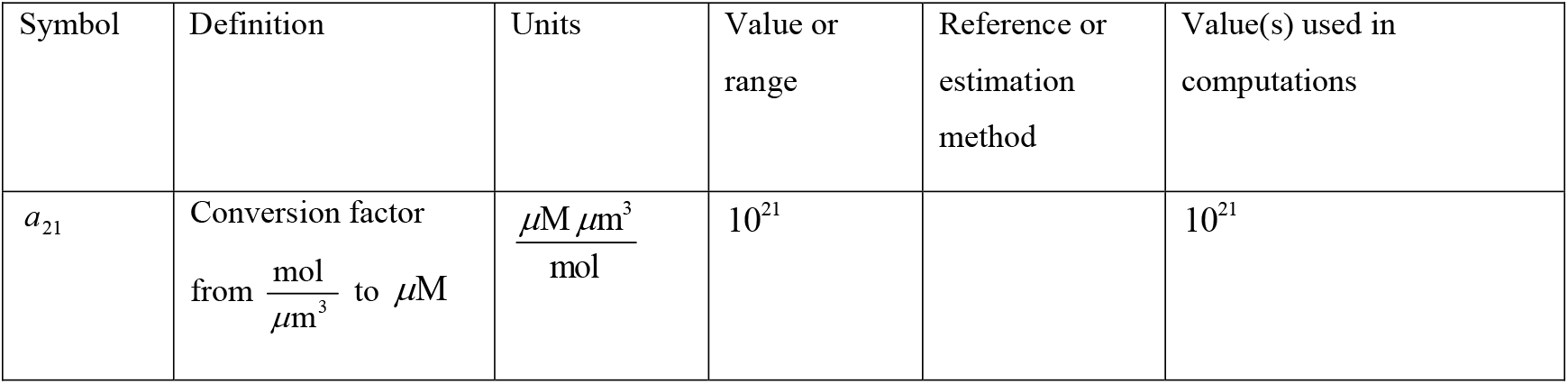
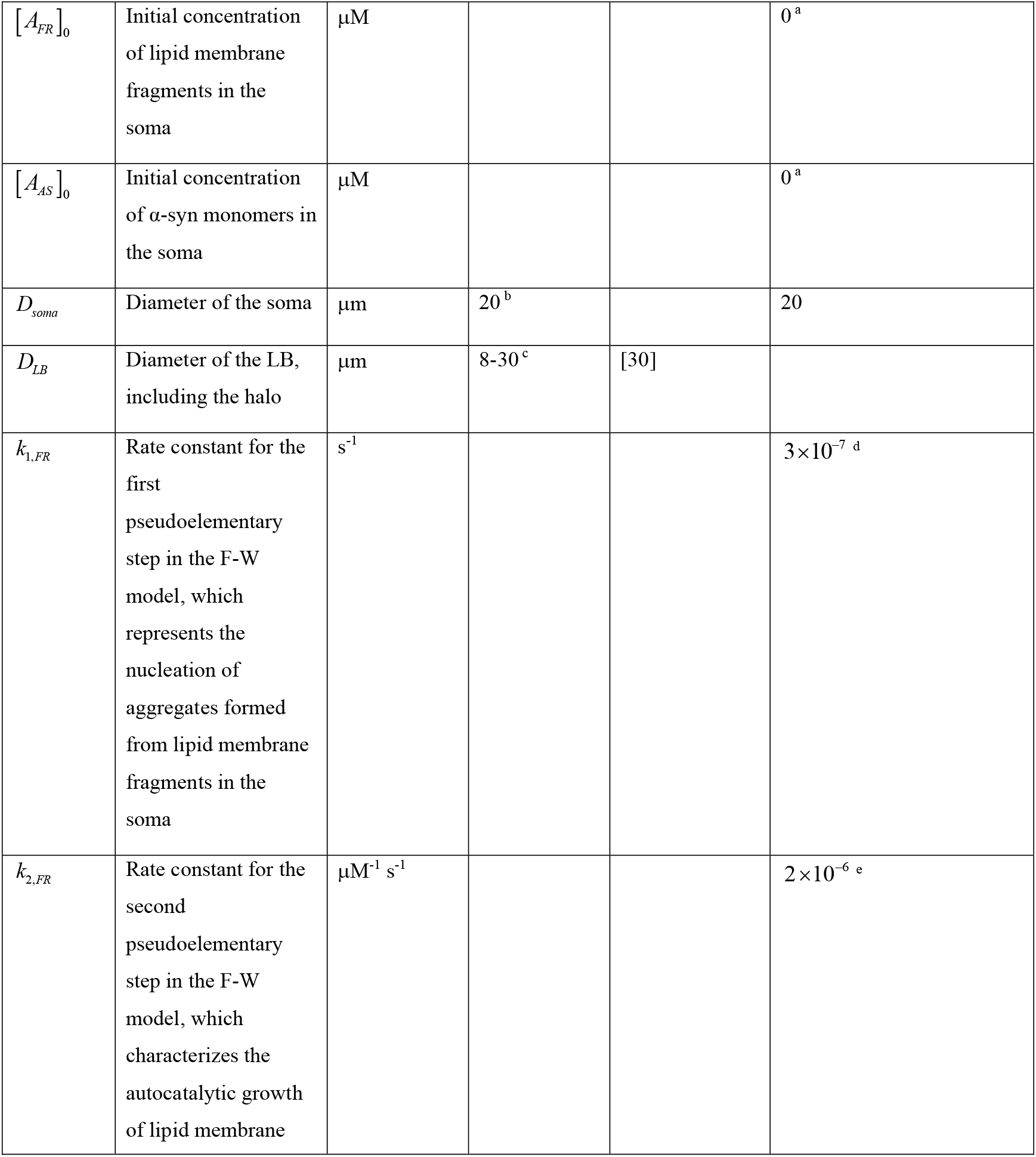
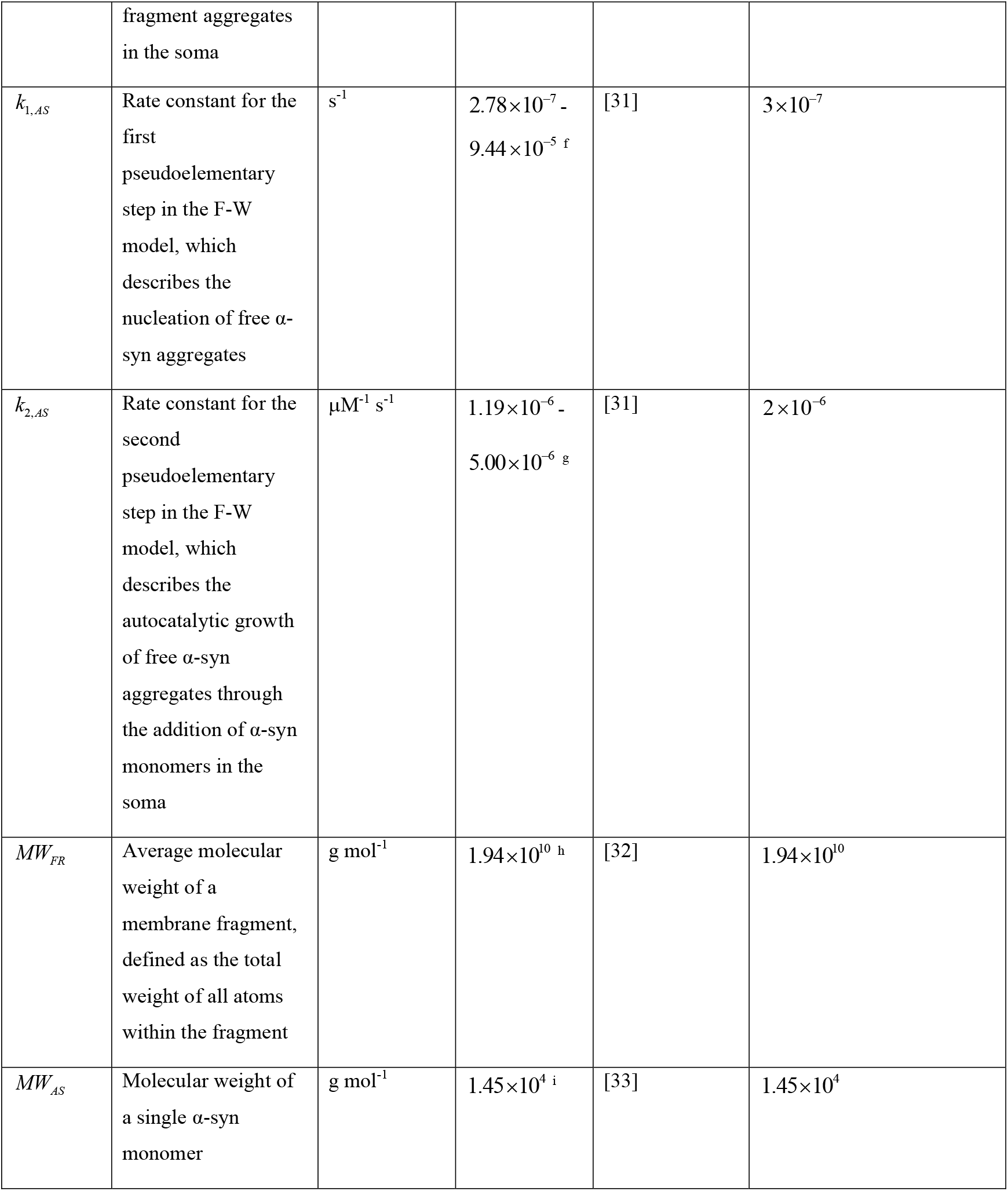
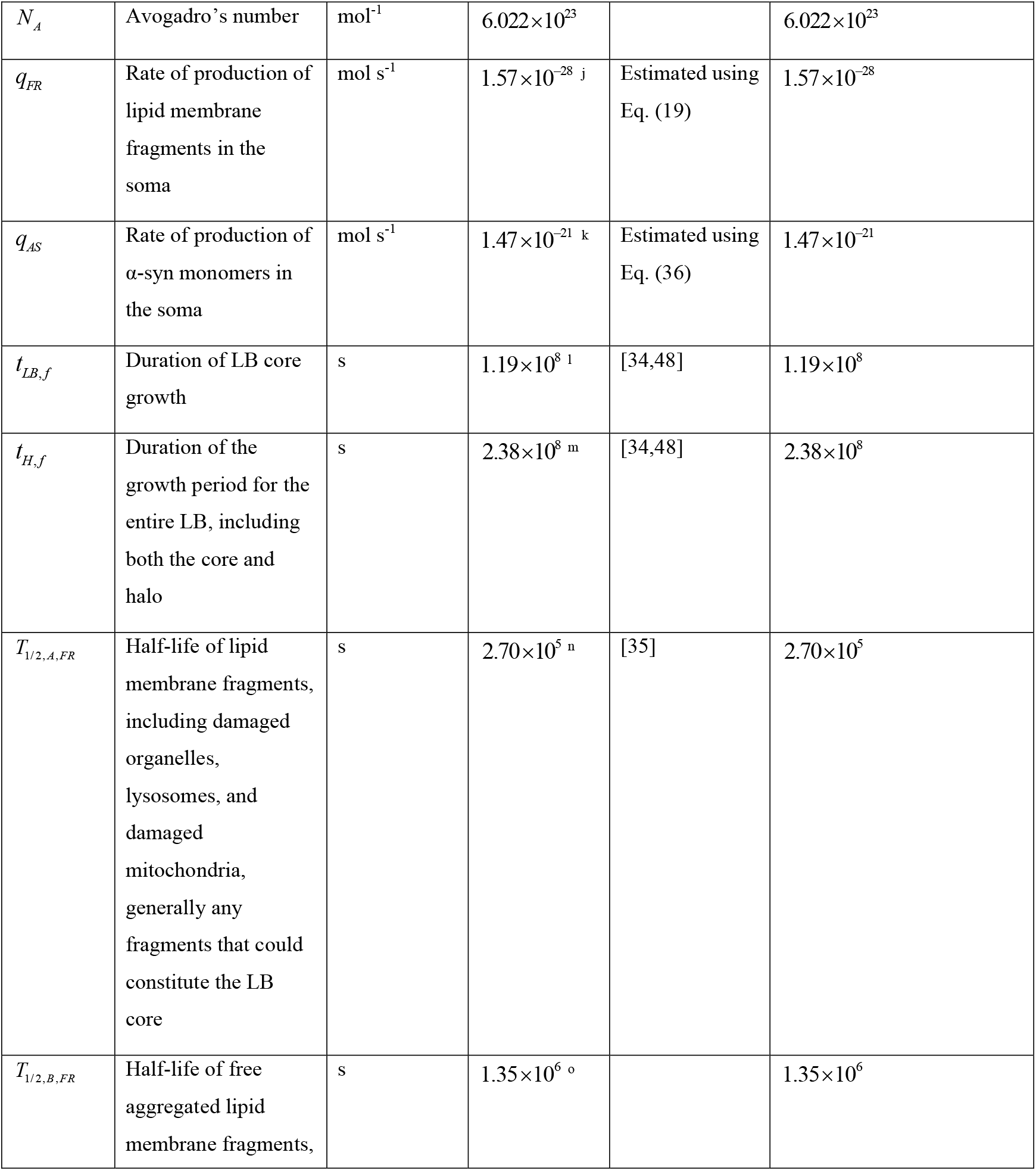
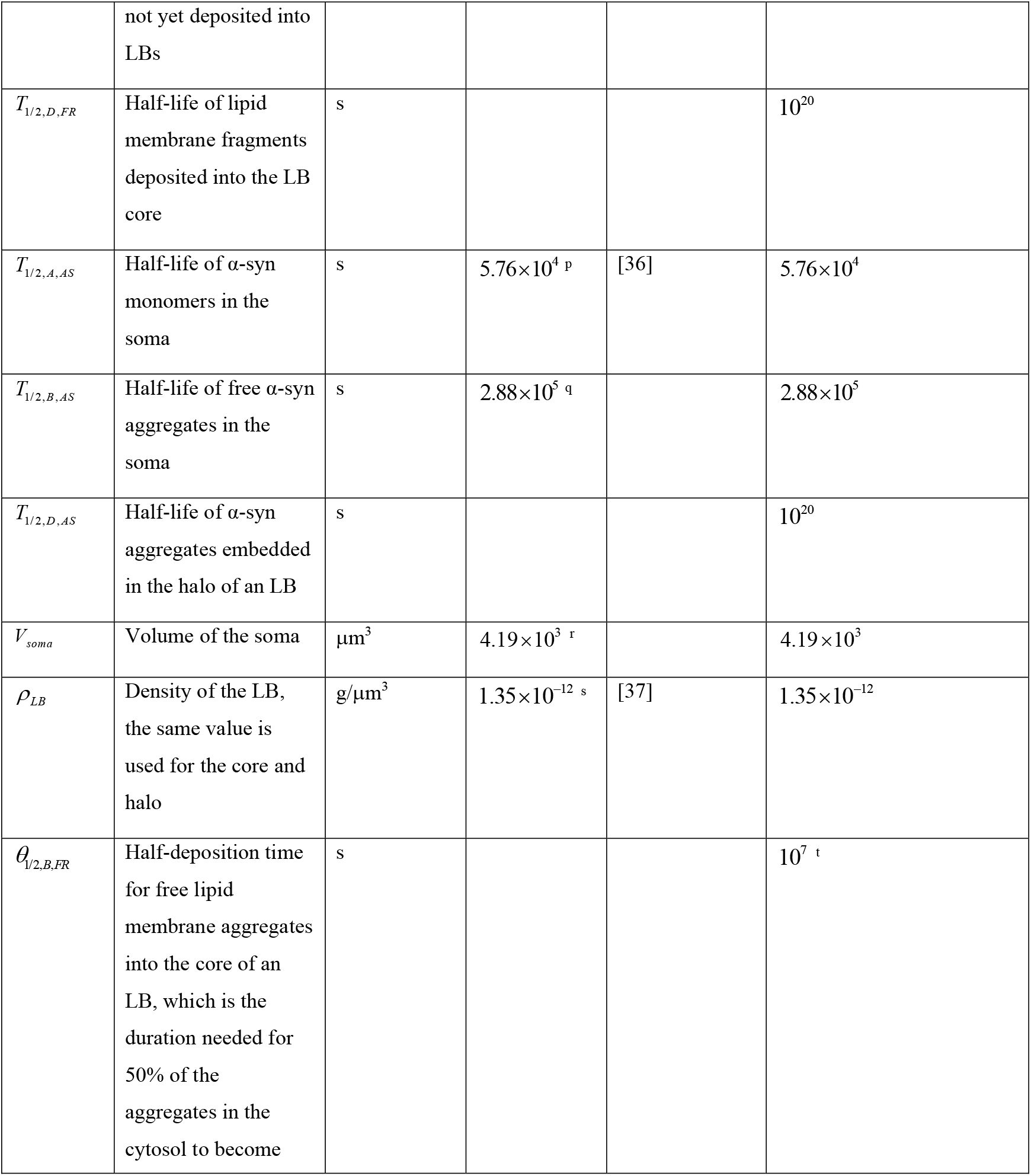
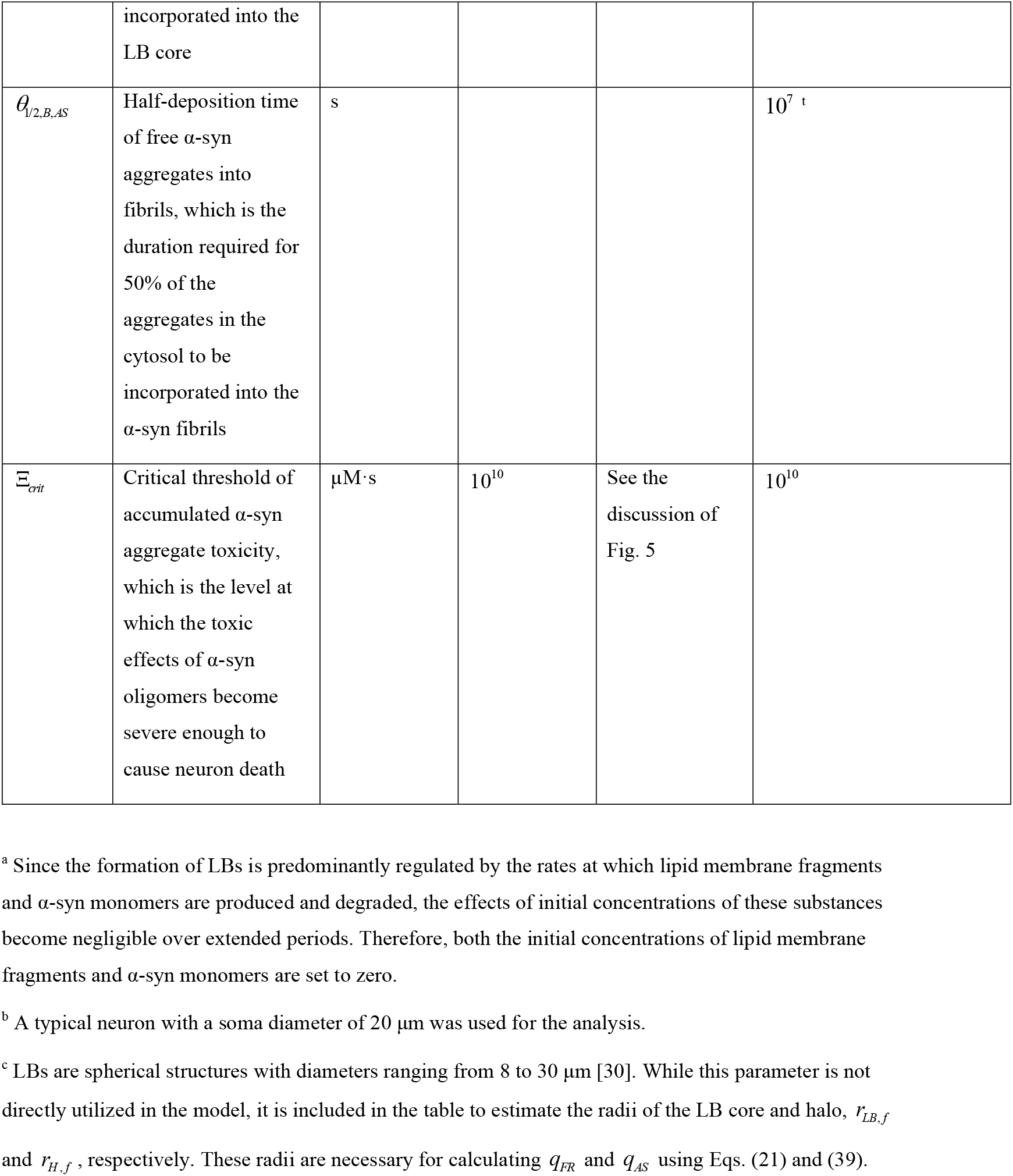
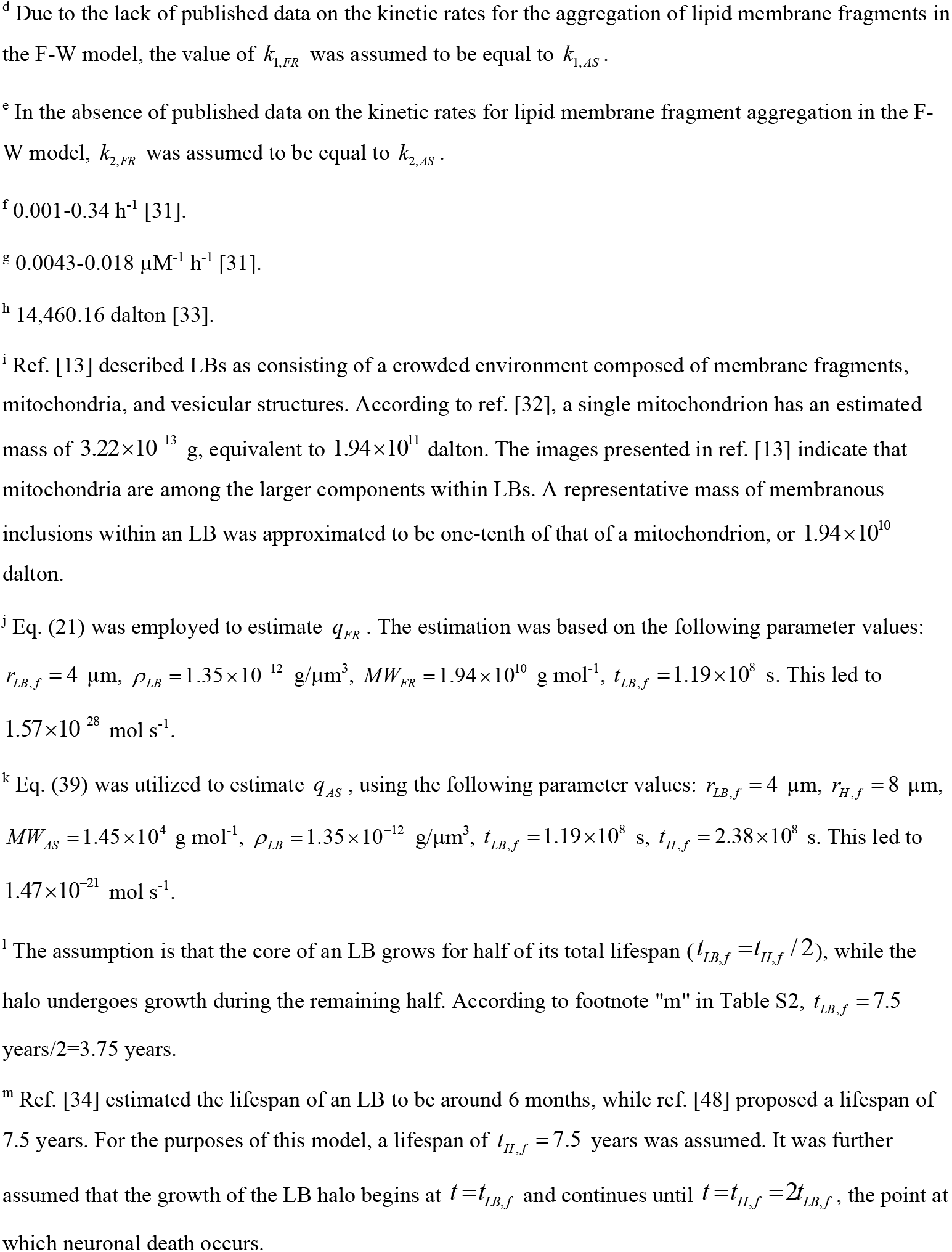
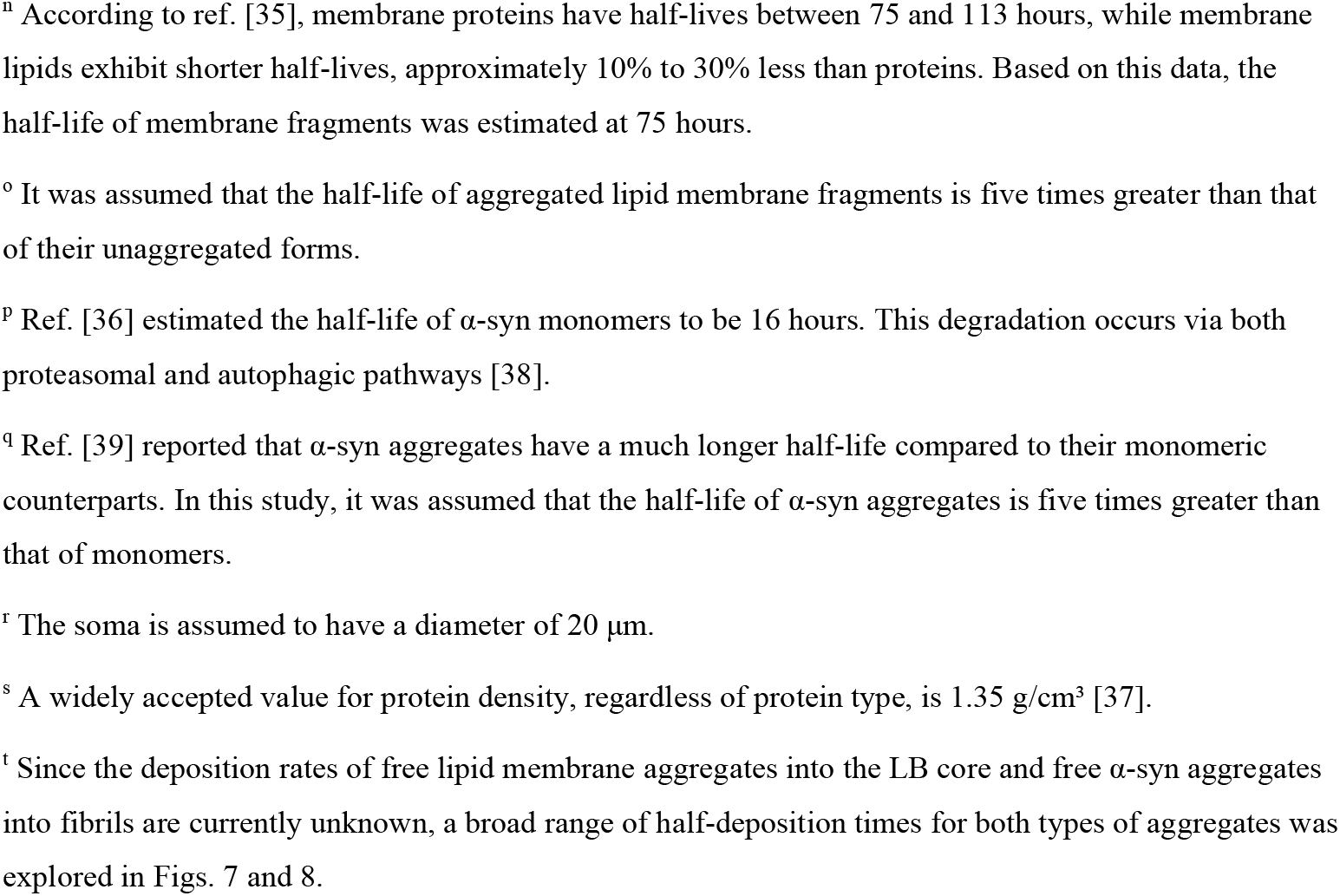
Model parameters and their estimated values.

| Symbol | Definition | Units | Value or range | Reference or estimation method | Value(s) used in computations |
| --- | --- | --- | --- | --- | --- |
| $a_{21}$ | Conversion factor from $\frac{\text{mol}}{\mu\text{m}^3}$ to $\mu\text{M}$ | $\frac{\mu\text{M } \mu\text{m}^3}{\text{mol}}$ | $10^{21}$ | | $10^{21}$ |
| $[A_{FR}]_0$ | Initial concentration of lipid membrane fragments in the soma | $\mu\text{M}$ | | | $0^a$ |
| $[A_{AS}]_0$ | Initial concentration of α-syn monomers in the soma | $\mu\text{M}$ | | | $0^a$ |
| $D_{soma}$ | Diameter of the soma | $\mu\text{m}$ | $20^b$ | | 20 |
| $D_{LB}$ | Diameter of the LB, including the halo | $\mu\text{m}$ | $8\text{-}30^c$ | [30] | |
| $k_{1,FR}$ | Rate constant for the first pseudoelementary step in the F-W model, which represents the nucleation of aggregates formed from lipid membrane fragments in the soma | $\text{s}^{-1}$ | | | $3 \times 10^{-7}^d$ |
| $k_{2,FR}$ | Rate constant for the second pseudoelementary step in the F-W model, which characterizes the autocatalytic growth of lipid membrane fragment aggregates in the soma | $\mu\text{M}^{-1} \text{s}^{-1}$ | | | $2 \times 10^{-6}^e$ |
| $k_{1,AS}$ | Rate constant for the first pseudoelementary step in the F-W model, which describes the nucleation of free α-syn aggregates | $s^{-1}$ | $2.78 \times 10^{-7}$ - $9.44 \times 10^{-5}$ f | [31] | $3 \times 10^{-7}$ |
| $k_{2,AS}$ | Rate constant for the second pseudoelementary step in the F-W model, which describes the autocatalytic growth of free α-syn aggregates through the addition of α-syn monomers in the soma | $\mu M^{-1} s^{-1}$ | $1.19 \times 10^{-6}$ - $5.00 \times 10^{-6}$ g | [31] | $2 \times 10^{-6}$ |
| $MW_{FR}$ | Average molecular weight of a membrane fragment, defined as the total weight of all atoms within the fragment | $g\ mol^{-1}$ | $1.94 \times 10^{10}$ h | [32] | $1.94 \times 10^{10}$ |
| $MW_{AS}$ | Molecular weight of a single α-syn monomer | $g\ mol^{-1}$ | $1.45 \times 10^4$ i | [33] | $1.45 \times 10^4$ |
| $N_A$ | Avogadro's number | $\text{mol}^{-1}$ | $6.022 \times 10^{23}$ | | $6.022 \times 10^{23}$ |
| $q_{FR}$ | Rate of production of lipid membrane fragments in the soma | $\text{mol s}^{-1}$ | $1.57 \times 10^{-28} \text{ j}$ | Estimated using Eq. (19) | $1.57 \times 10^{-28}$ |
| $q_{AS}$ | Rate of production of α-syn monomers in the soma | $\text{mol s}^{-1}$ | $1.47 \times 10^{-21} \text{ k}$ | Estimated using Eq. (36) | $1.47 \times 10^{-21}$ |
| $t_{LB,f}$ | Duration of LB core growth | s | $1.19 \times 10^8 \text{ l}$ | [34,48] | $1.19 \times 10^8$ |
| $t_{H,f}$ | Duration of the growth period for the entire LB, including both the core and halo | s | $2.38 \times 10^8 \text{ m}$ | [34,48] | $2.38 \times 10^8$ |
| $T_{1/2,A,FR}$ | Half-life of lipid membrane fragments, including damaged organelles, lysosomes, and damaged mitochondria, generally any fragments that could constitute the LB core | s | $2.70 \times 10^5 \text{ n}$ | [35] | $2.70 \times 10^5$ |
| $T_{1/2,B,FR}$ | Half-life of free aggregated lipid membrane fragments, not yet deposited into LBs | s | $1.35 \times 10^6 \text{ o}$ | | $1.35 \times 10^6$ |
| $T_{1/2,D,FR}$ | Half-life of lipid membrane fragments deposited into the LB core | s | | | $10^{20}$ |
| $T_{1/2,A,AS}$ | Half-life of α-syn monomers in the soma | s | $5.76 \times 10^4$ p | [36] | $5.76 \times 10^4$ |
| $T_{1/2,B,AS}$ | Half-life of free α-syn aggregates in the soma | s | $2.88 \times 10^5$ q | | $2.88 \times 10^5$ |
| $T_{1/2,D,AS}$ | Half-life of α-syn aggregates embedded in the halo of an LB | s | | | $10^{20}$ |
| $V_{soma}$ | Volume of the soma | $\mu\text{m}^3$ | $4.19 \times 10^3$ r | | $4.19 \times 10^3$ |
| $\rho_{LB}$ | Density of the LB, the same value is used for the core and halo | $\text{g}/\mu\text{m}^3$ | $1.35 \times 10^{-12}$ s | [37] | $1.35 \times 10^{-12}$ |
| $\theta_{1/2,B,FR}$ | Half-deposition time for free lipid membrane aggregates into the core of an LB, which is the duration needed for 50% of the aggregates in the cytosol to become incorporated into the LB core | s | | | $10^7$ t |
| $\theta_{1/2,B,AS}$ | Half-deposition time of free α-syn aggregates into fibrils, which is the duration required for 50% of the aggregates in the cytosol to be incorporated into the α-syn fibrils | s | | | $10^7$ <sup>t</sup> |
| $\Xi_{crit}$ | Critical threshold of accumulated α-syn aggregate toxicity, which is the level at which the toxic effects of α-syn oligomers become severe enough to cause neuron death | $\mu\text{M}\cdot\text{s}$ | $10^{10}$ | See the discussion of Fig. 5 | $10^{10}$ |

<sup>a</sup> Since the formation of LBs is predominantly regulated by the rates at which lipid membrane fragments and α-syn monomers are produced and degraded, the effects of initial concentrations of these substances become negligible over extended periods. Therefore, both the initial concentrations of lipid membrane fragments and α-syn monomers are set to zero.
<sup>b</sup> A typical neuron with a soma diameter of 20 $\mu\text{m}$ was used for the analysis.
<sup>c</sup> LBs are spherical structures with diameters ranging from 8 to 30 $\mu\text{m}$ [30]. While this parameter is not directly utilized in the model, it is included in the table to estimate the radii of the LB core and halo, $r_{LB,f}$ and $r_{H,f}$ , respectively. These radii are necessary for calculating $q_{FR}$ and $q_{AS}$ using Eqs. (21) and (39).
<sup>d</sup> Due to the lack of published data on the kinetic rates for the aggregation of lipid membrane fragments in the F-W model, the value of $k_{1,FR}$ was assumed to be equal to $k_{1,AS}$ .
<sup>e</sup> In the absence of published data on the kinetic rates for lipid membrane fragment aggregation in the F-W model, $k_{2,FR}$ was assumed to be equal to $k_{2,AS}$ .
<sup>f</sup> 0.001-0.34 h<sup>-1</sup> [31].
<sup>g</sup> 0.0043-0.018 μM<sup>-1</sup> h<sup>-1</sup> [31].
<sup>h</sup> 14,460.16 dalton [33].
<sup>i</sup> Ref. [13] described LBs as consisting of a crowded environment composed of membrane fragments, mitochondria, and vesicular structures. According to ref. [32], a single mitochondrion has an estimated mass of $3.22 \times 10^{-13}$ g, equivalent to $1.94 \times 10^{11}$ dalton. The images presented in ref. [13] indicate that mitochondria are among the larger components within LBs. A representative mass of membranous inclusions within an LB was approximated to be one-tenth of that of a mitochondrion, or $1.94 \times 10^{10}$ dalton.
<sup>j</sup> Eq. (21) was employed to estimate $q_{FR}$ . The estimation was based on the following parameter values:
$r_{LB,f} = 4$ μm, $\rho_{LB} = 1.35 \times 10^{-12}$ g/μm<sup>3</sup>, $MW_{FR} = 1.94 \times 10^{10}$ g mol<sup>-1</sup>, $t_{LB,f} = 1.19 \times 10^8$ s. This led to $1.57 \times 10^{-28}$ mol s<sup>-1</sup>.
<sup>k</sup> Eq. (39) was utilized to estimate $q_{AS}$ , using the following parameter values: $r_{LB,f} = 4$ μm, $r_{H,f} = 8$ μm, $MW_{AS} = 1.45 \times 10^4$ g mol<sup>-1</sup>, $\rho_{LB} = 1.35 \times 10^{-12}$ g/μm<sup>3</sup>, $t_{LB,f} = 1.19 \times 10^8$ s, $t_{H,f} = 2.38 \times 10^8$ s. This led to $1.47 \times 10^{-21}$ mol s<sup>-1</sup>.
<sup>l</sup> The assumption is that the core of an LB grows for half of its total lifespan ( $t_{LB,f} = t_{H,f} / 2$ ), while the halo undergoes growth during the remaining half. According to footnote "m" in Table S2, $t_{LB,f} = 7.5$ years/2 = 3.75 years.
<sup>m</sup> Ref. [34] estimated the lifespan of an LB to be around 6 months, while ref. [48] proposed a lifespan of 7.5 years. For the purposes of this model, a lifespan of $t_{H,f} = 7.5$ years was assumed. It was further assumed that the growth of the LB halo begins at $t = t_{LB,f}$ and continues until $t = t_{H,f} = 2t_{LB,f}$ , the point at which neuronal death occurs.

### S4. Supplemental figures

**Fig. S1.**
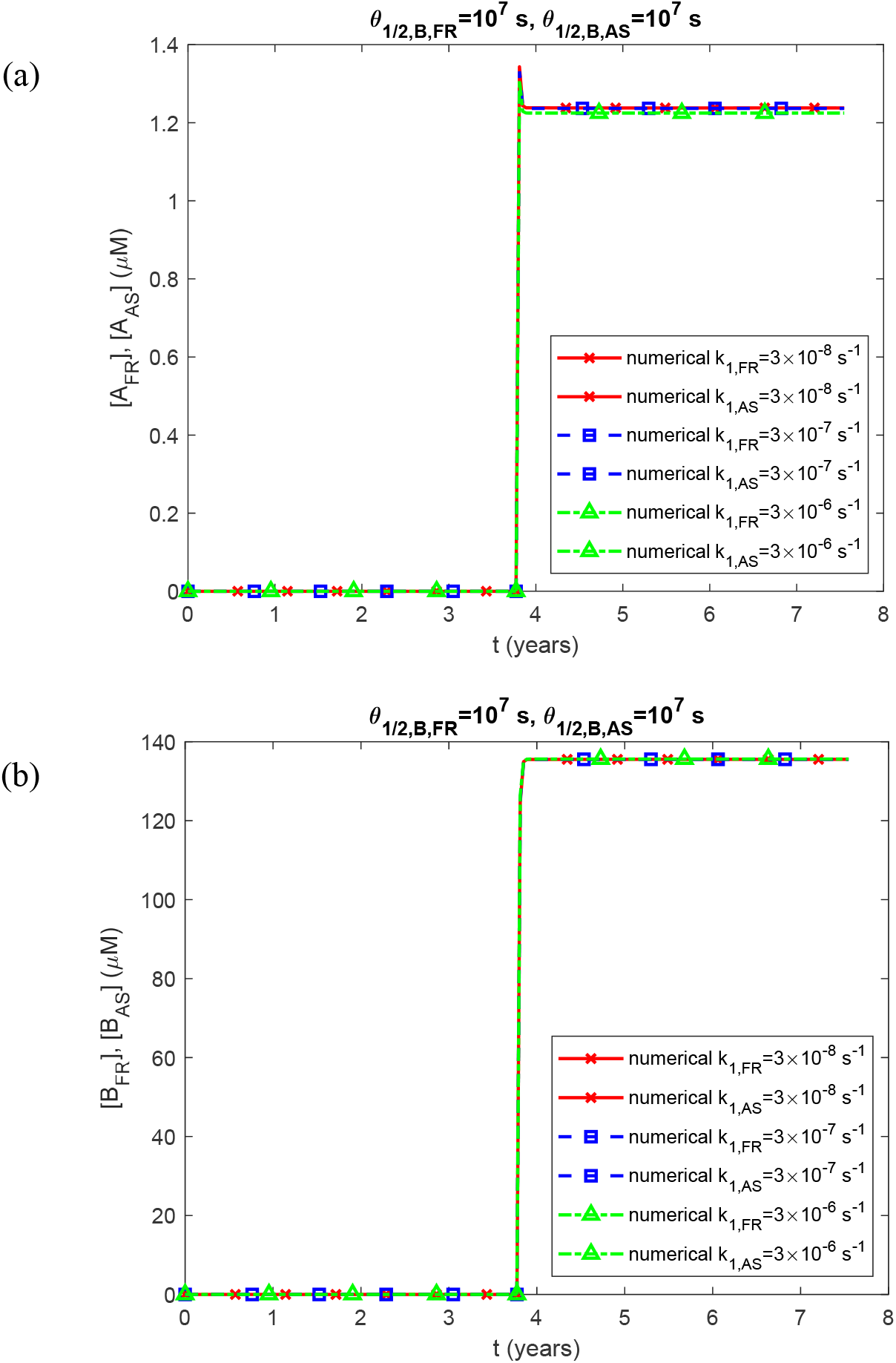
(a) Molar concentrations of lipid membrane fragments and α-syn monomers, [*A*_*FR*_] and [*A*_*AS*_], respectively, for different values of *k*_1, *FR*_ and *k*_1, *AS*_ vs time. (b) Molar concentrations of free lipid membrane aggregates and free α-syn aggregates, [*B*_*FR*_] and [*B*_*AS*_], respectively, for different values of *k* _1, *FR*_ and *k*_1, *AS*_ vs time. (*k*_2,*FR*_ = *k*_2, *AS*_ = 2×10^−6^ μM^−1^ s^−1^, *q* _*FR*_ =1.57×10^−28^ mol s^−1^, *q*_*AS*_ = 1.47 ×10^−21^ mol s^−1^.)

**Fig. S2.**
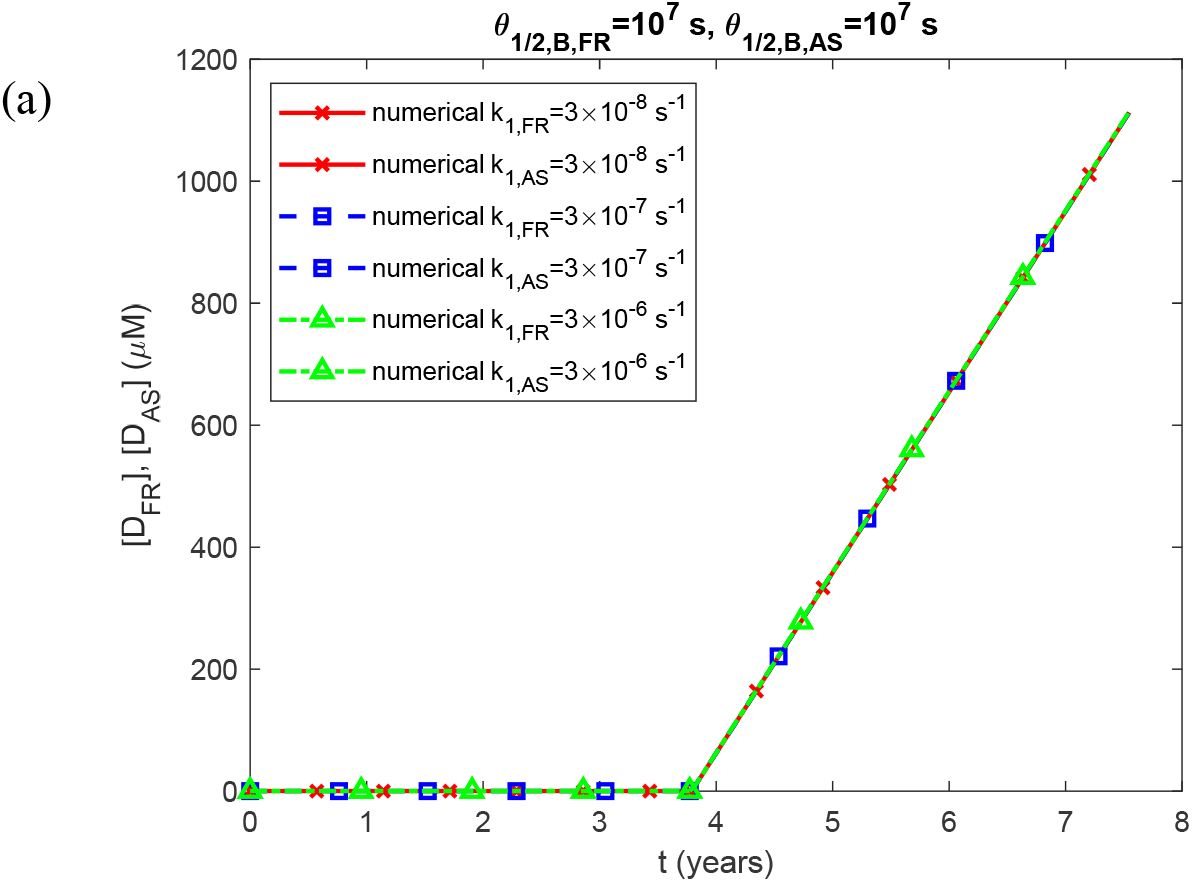
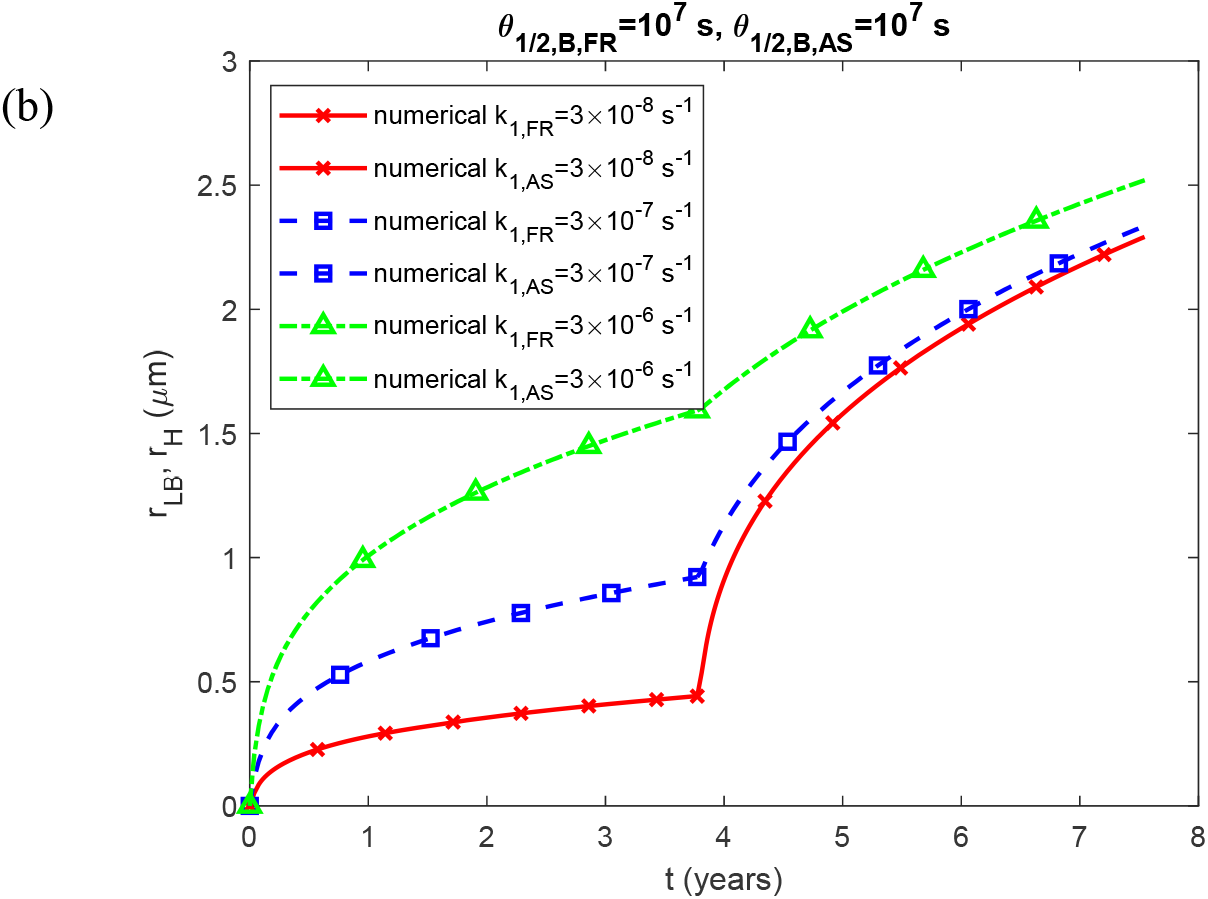
(a) Molar concentrations of lipid membrane aggregates deposited into the core of the LB and α-syn aggregates deposited in the halo’s fibrils, [*D*_*FR*_] and [*D*_*AS*_], respectively, for different values of *k*_1,*FR*_ and *k*_1, *AS*_ vs time. (b) Radii of the growing core of the LB and the growing halo, *r*_*LB*_ and *r*_*H*_, respectively, for different values of *k*_1, *FR*_ and *k*_1, *AS*_ vs time. (*k*_2,*FR*_ = *k*_2, *AS*_ = 2×10^−6^ μM^−1^ s^−1^, *q*_*FR*_ =1.57×10^−28^ mol s^−1^, *q* = 1.47 ×10^−21^ mol s^−1^.)

**Fig. S3.**
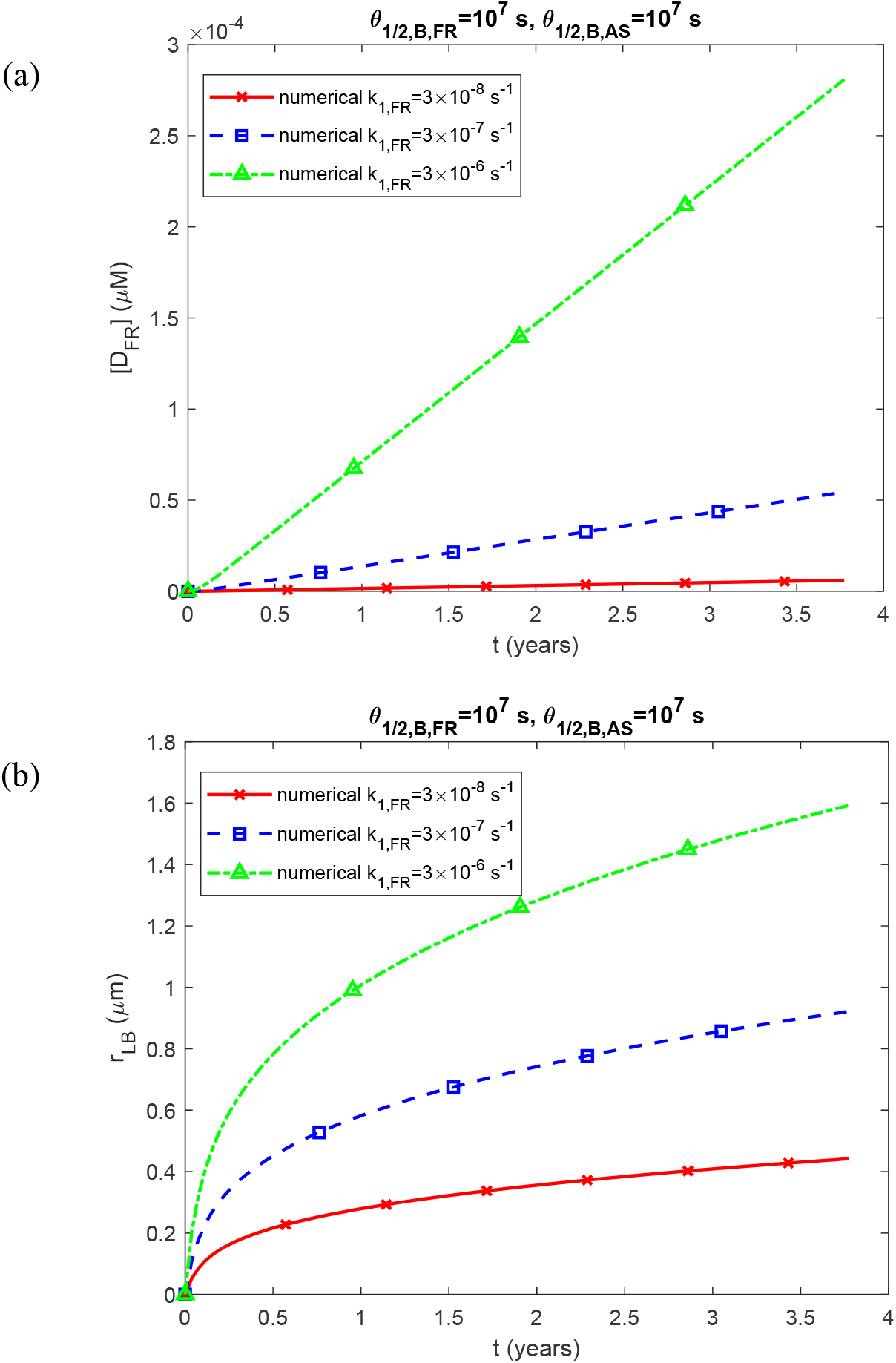
(a) Molar concentration of lipid membrane aggregates deposited into the core of the LB, [*D*_*FR*_], for different values of *k*_1, *FR*_ and *k*_1, *AS*_ vs time. (b) Radius of the growing core of the LB, *r*_*LB*_, for different values of *k*_1, *FR*_ and *k*_1, *AS*_ vs time. (*k*_2,*FR*_ = *k*_2, *AS*_ = 2×10^−6^ μM^−1^ s^−1^, *q*_*FR*_ =1.57×10^−28^ mol s^−1^, *q*_*AS*_ = 1.47 ×10^−21^ mol s^−1^.)

**Fig. S4.**
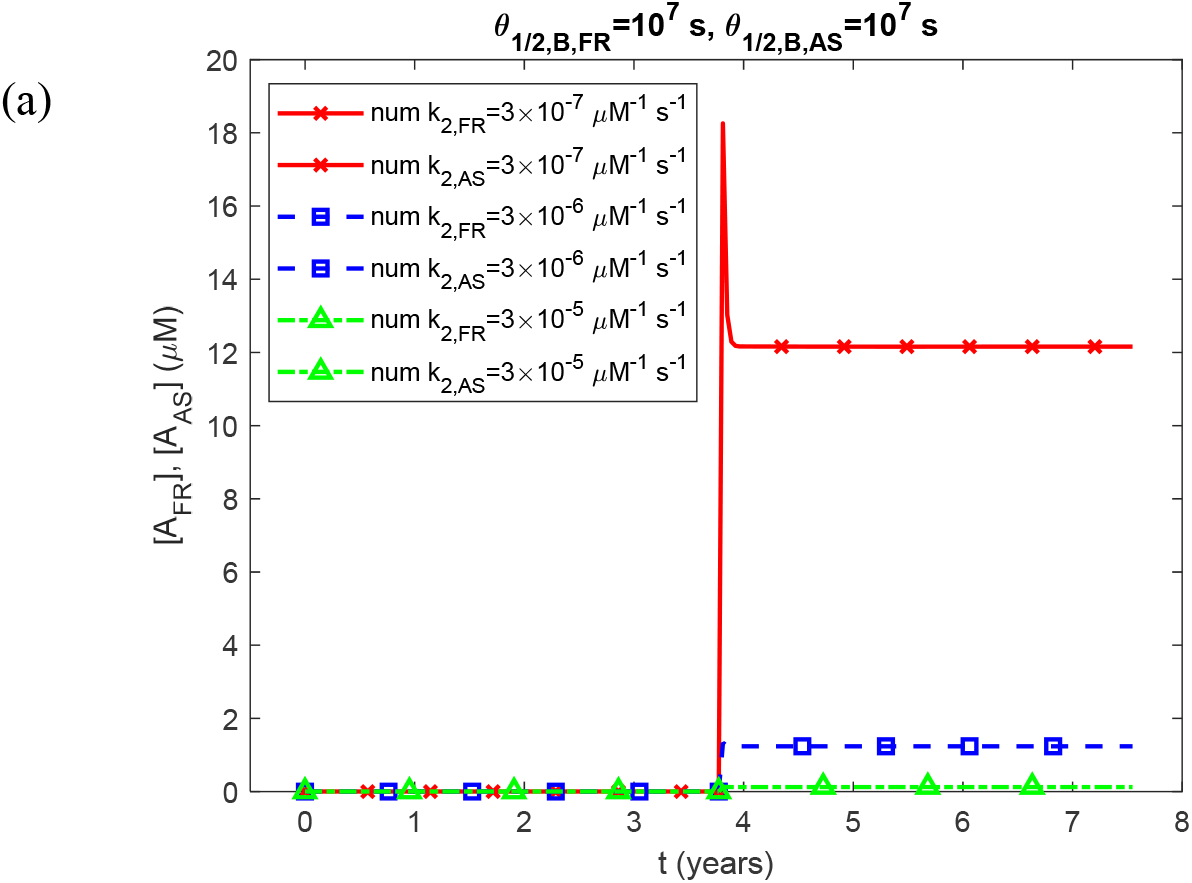
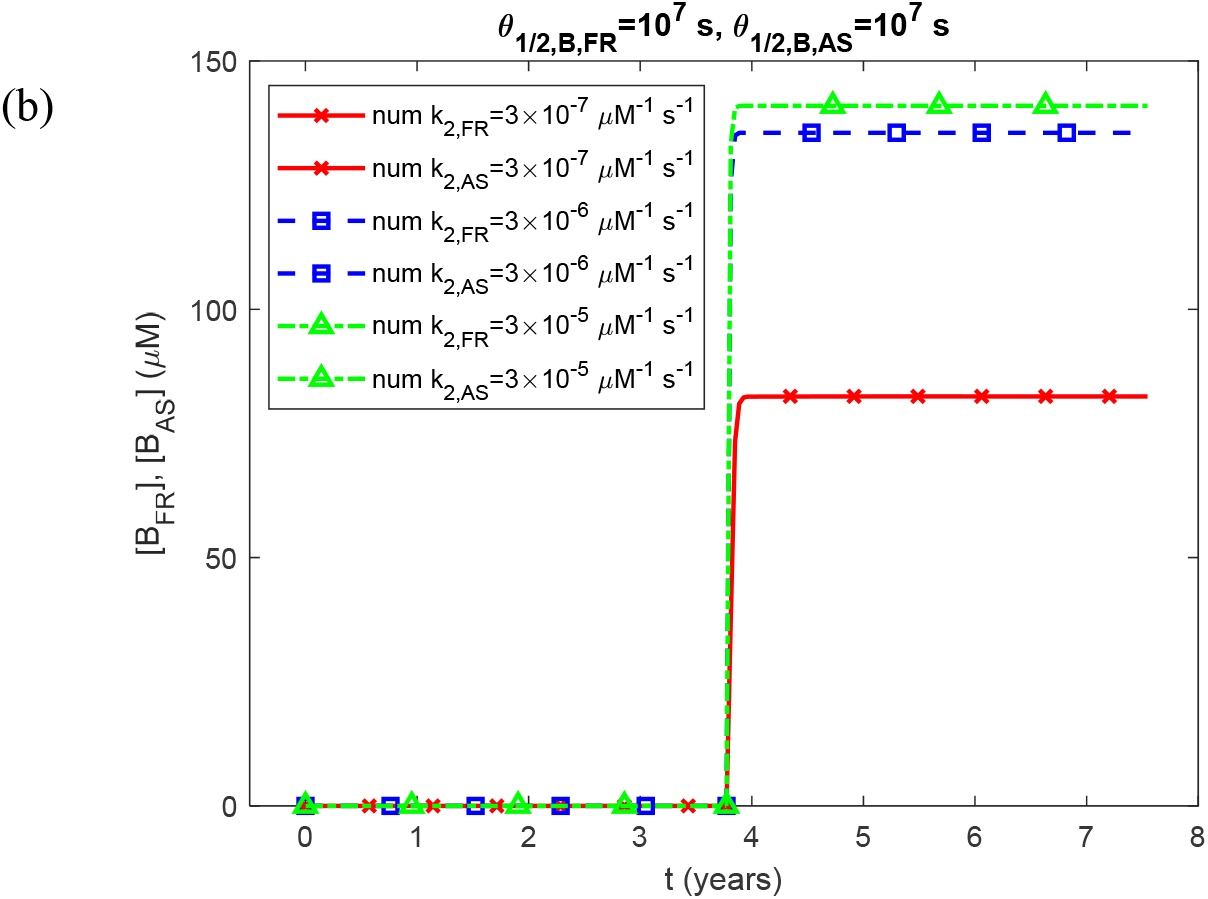
(a) Molar concentrations of lipid membrane fragments and α-syn monomers, [*A*_*FR*_] and [*A*_*AS*_], respectively, for different values of *k*_2, *FR*_ and *k*_2, *AS*_ vs time. (b) Molar concentrations of free lipid membrane aggregates and free α-syn aggregates, [*B*_*FR*_] and [*B*_*AS*_], respectively, for different values of *k*_2, *FR*_ and *k*_2, *AS*_ vs time. (*k*_1,*FR*_ = *k*_1, *AS*_ = 3×10^−7^ s^−1^, *q*_*FR*_ =1.57×10^−28^ mol s^−1^, *q*_*AS*_ = 1.47 ×10^−21^ mol s^−1^.) In the legend, “num” stands for “numerical.”

**Fig. S5.**
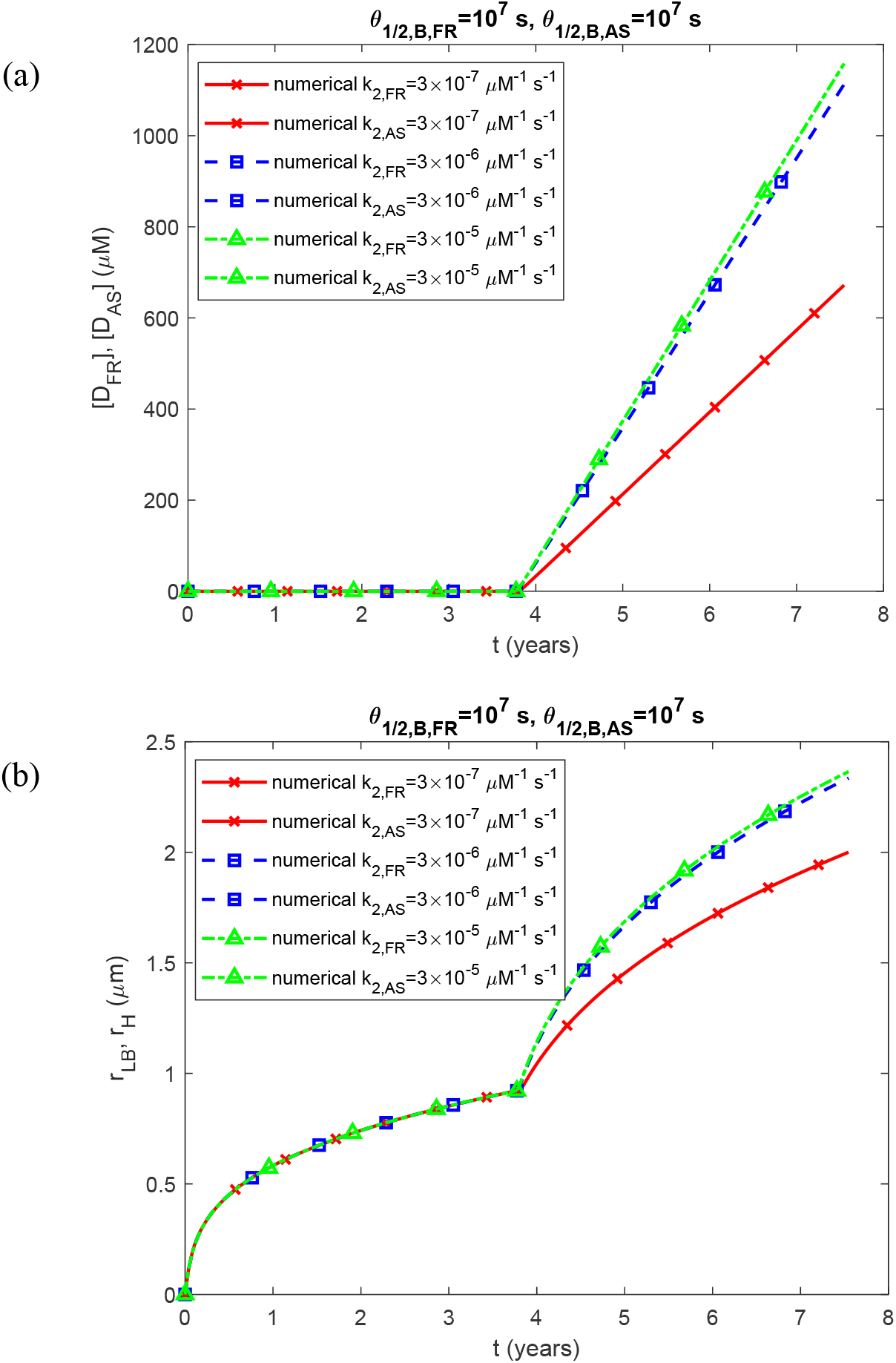
(a) Molar concentrations of lipid membrane aggregates deposited into the core of the LB and α-syn aggregates deposited in the halo’s fibrils, [*D*_*FR*_] and [*D*_*AS*_], respectively, for different values of *k*_2,*FR*_ and *k*_2, *AS*_ vs time. (b) Radii of the growing core of the LB and the growing halo, *r*_*LB*_ and *r*_*H*_, respectively, for different values of *k*_2, *FR*_ and *k*_2, *AS*_ vs time. (*k*_1,*FR*_ = *k*_1, *AS*_ = 3×10^−7^ s^−1^, *q*_*FR*_ =1.57×10^−28^ mol s^−1^, *q*_*AS*_ = 1.47 ×10^−21^ mol s^−1^.)

**Fig. S6.**
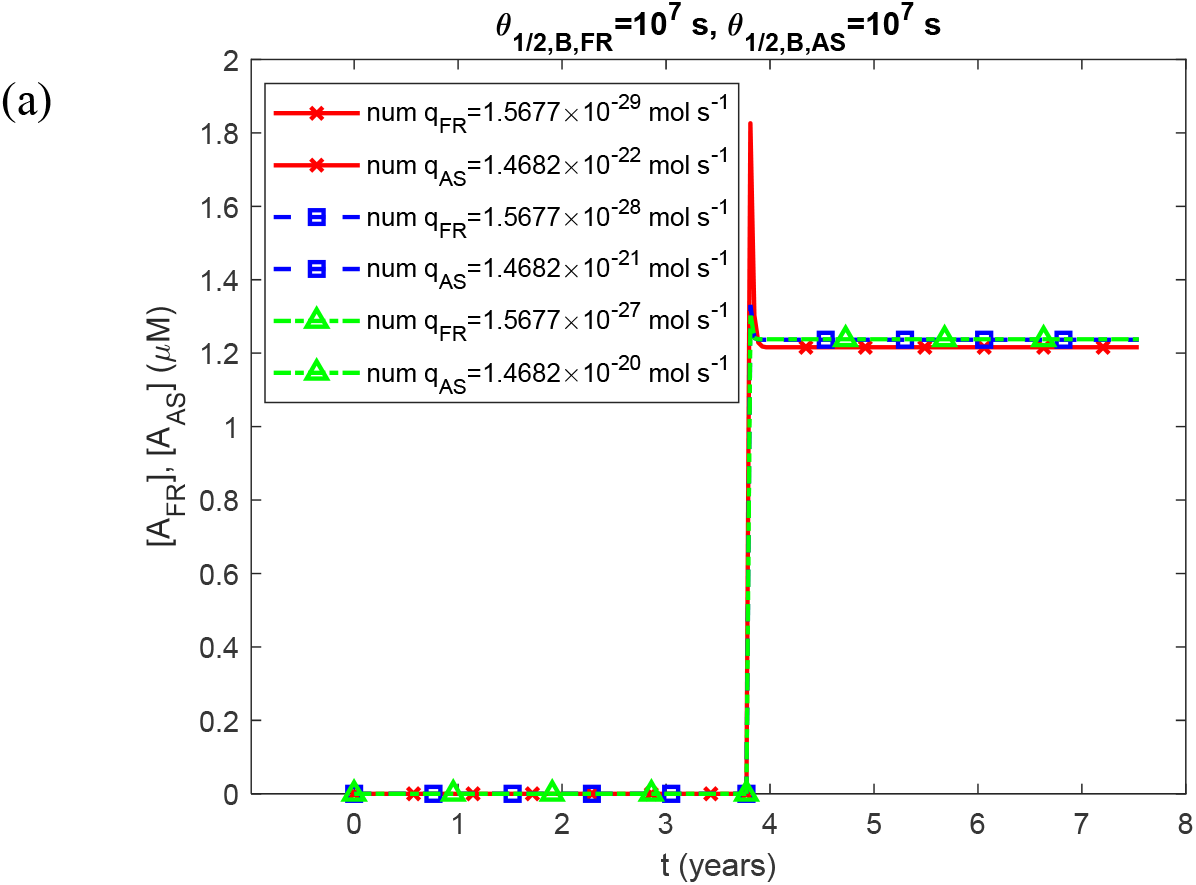
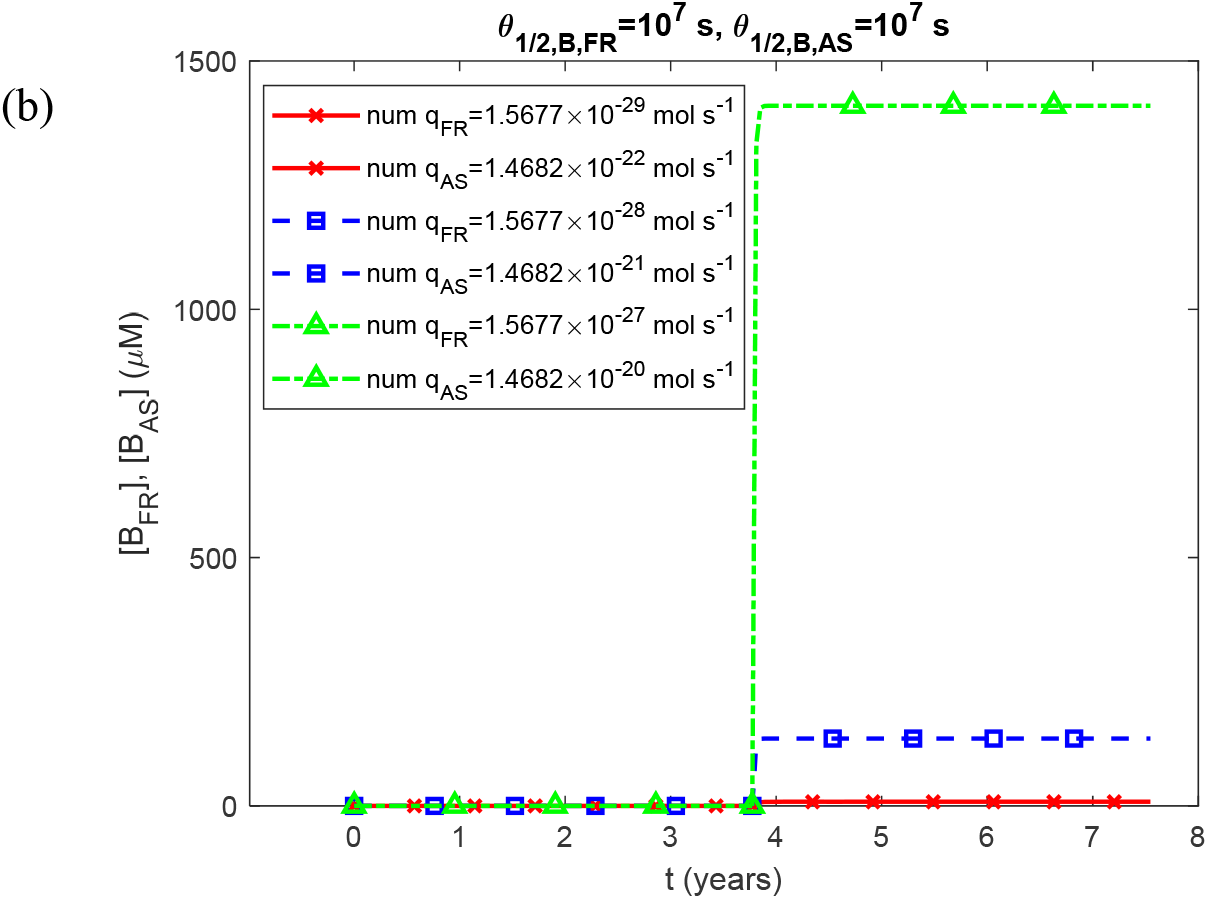
(a) Molar concentrations of lipid membrane fragments and α-syn monomers, [*A*_*FR*_] and [*A* _*AS*_] respectively, for different values of *q*_*FR*_ and *q*_*AS*_ vs time. (b) Molar concentrations of free lipid membrane aggregates and free α-syn aggregates, [*B*_*FR*_] and [*B*_*AS*_], respectively, for different values of *q*_*FR*_ and *q*_*AS*_ vs time. (*k*_1,*FR*_ = *k*_1, *AS*_ = 3×10^−7^ s^−1^, *k*_2, *FR*_ = *k*_2, *AS*_ = 2×10^−6^ μM^−1^ s^−1^.) In the legend, “num” stands for “numerical.”

**Fig. S7.**
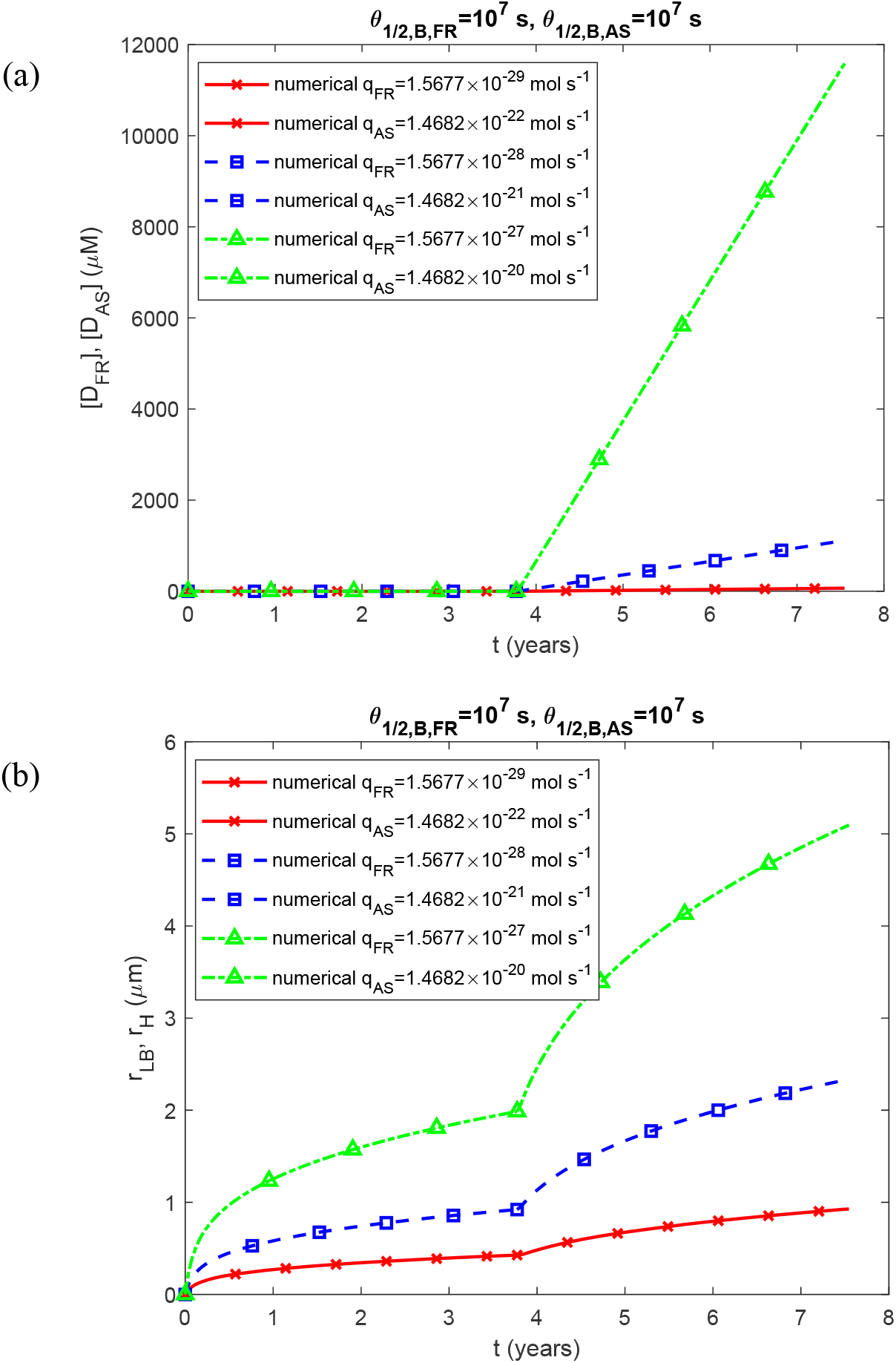
(a) Molar concentrations of lipid membrane aggregates deposited into the core of the LB and α-syn aggregates deposited in the halo’s fibrils, [*D*_*FR*_] and [*D*_*AS*_], respectively, for different values of *q*_*FR*_ and *q*_*AS*_ vs time. (b) Radii of the growing core of the LB and the growing halo, *r*_*LB*_ and *r*_*H*_, respectively, for different values of *q*_*FR*_ and *q*_*AS*_ vs time. (*k*_1,*FR*_ = *k*_1, *AS*_ = 3×10^−7^ s^−1^, *k*_2, *FR*_ = *k*_2, *AS*_ = 2×10^−6^ μM^−1^ s^−1^.)

**Fig. S8.**
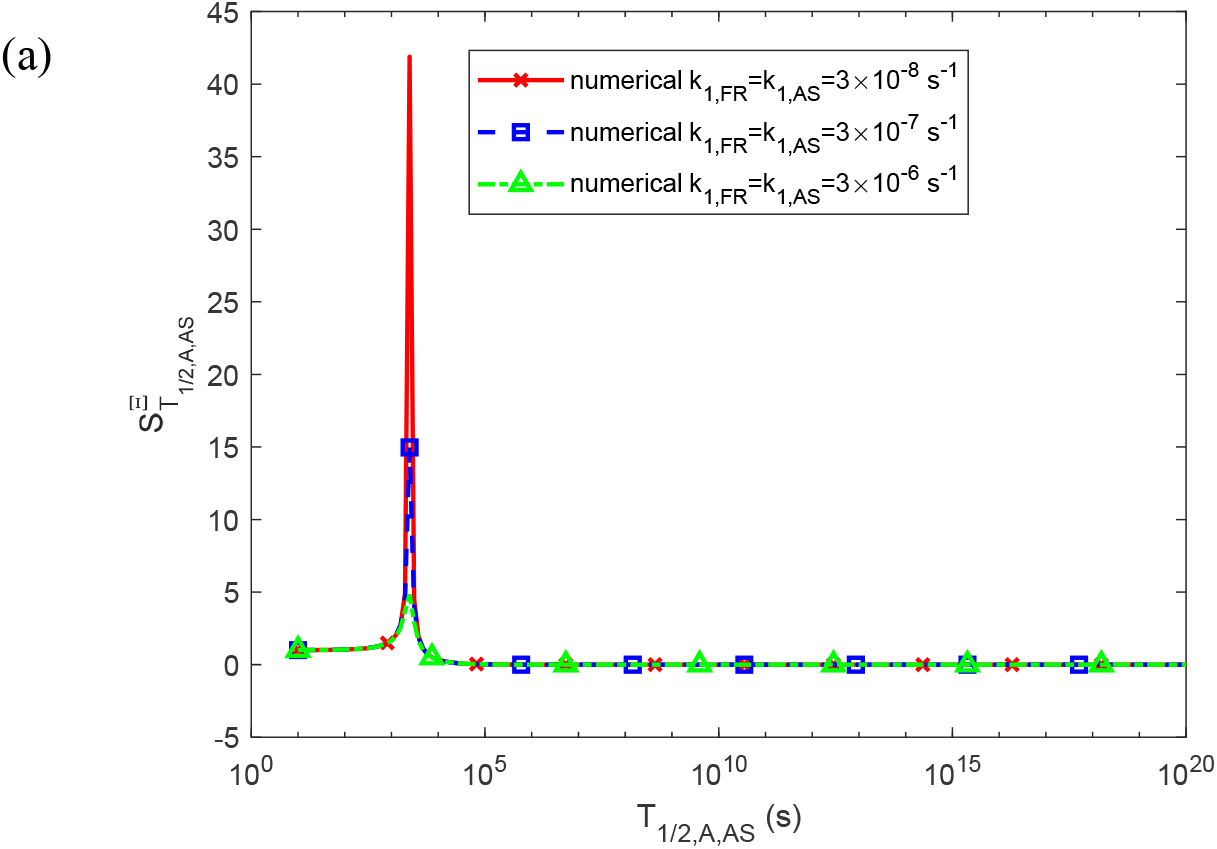
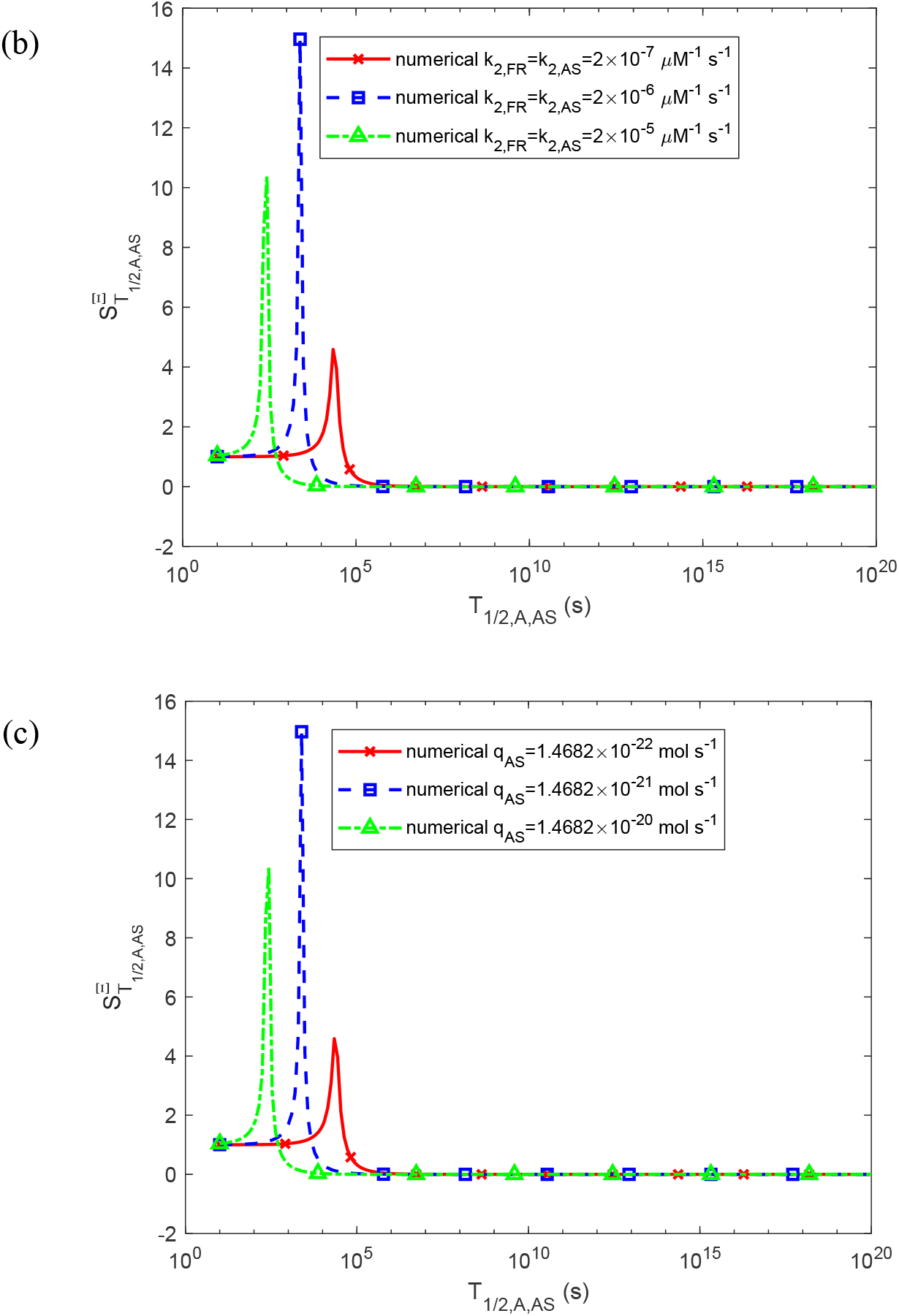
Sensitivity of accumulated toxicity, Ξ, to the half-life of α-syn monomers, *T*_1/ 2, *A, AS*_, is examined for three different values of (a) the rate constants that describes the nucleation of membrane fragments and α-syn aggregates, *k*_1,*FR*_ and *k*_1, *AS*_ (*k*_2, *FR*_ = *k*_2, *AS*_ = 2×10^−6^ μM^−1^ s^−1^, *q*_*FR*_ =1.57×10^−28^ mol s^−1^, *q*_*AS*_ = 1.47 ×10^−21^ mol s^−1^), (b) the rate constants that describe the autocatalytic growth of membrane fragments and α-syn aggregates, *k*_2, *FR*_ and *k*_2, *AS*_ (*k*_1,*FR*_ = *k*_1, *AS*_ = 3×10^−7^ s^−1^, *q*_*FR*_ =1.57×10^−28^ mol s^−1^, *q*_*AS*_ = 1.47 ×10^−21^ mol s^−1^), (c) the production rate of α-syn monomers, *q*_*AS*_. (*k*_1,*FR*_ = *k*_1, *AS*_ = 3×10^−7^ s^−1^, *k*_2, *FR*_ = *k*_2, *AS*_ = 2×10^−6^ μM^−1^ s^−1^). _1/ 2, *B, AS*_ *T*_1/ 2, *A, FR*_, *T*_1/ 2, *B, FR*_, and *T* were kept at their values given in Table S2.

**Fig. S9.**
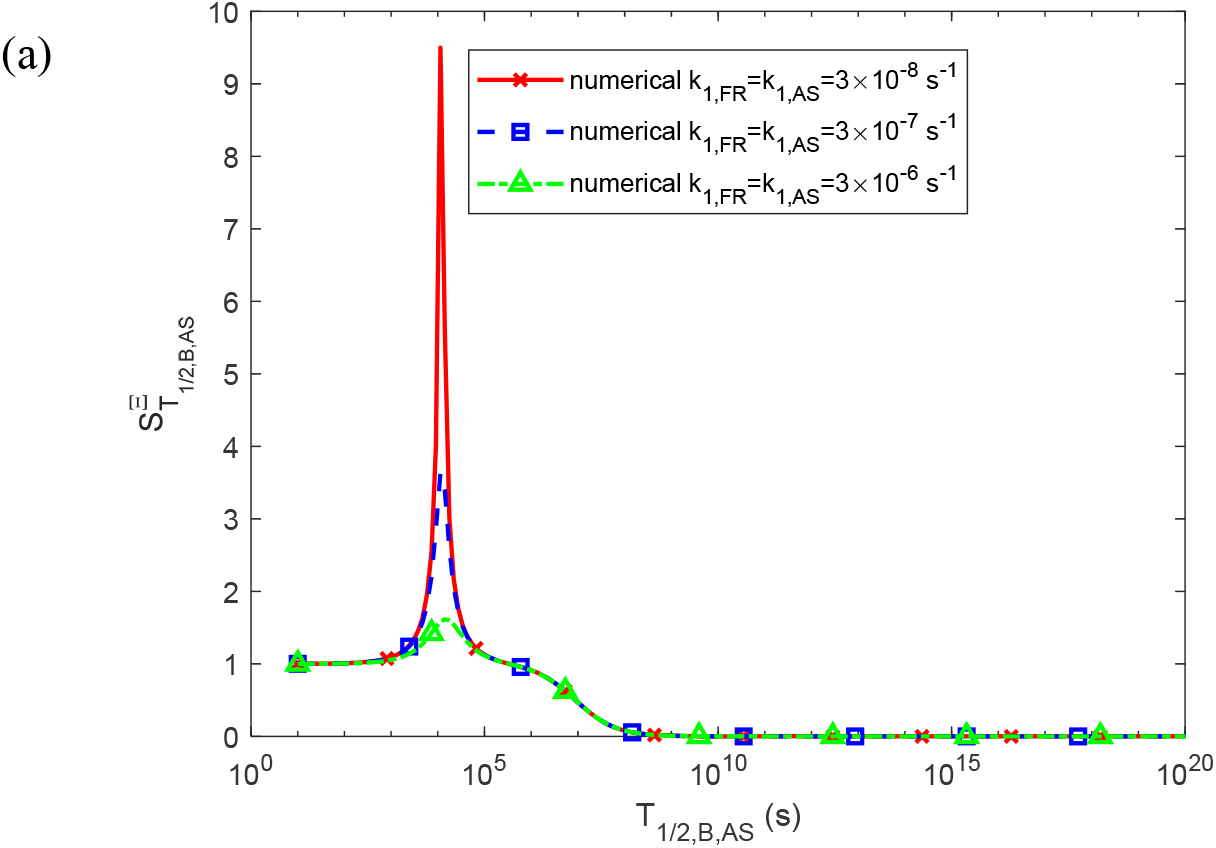
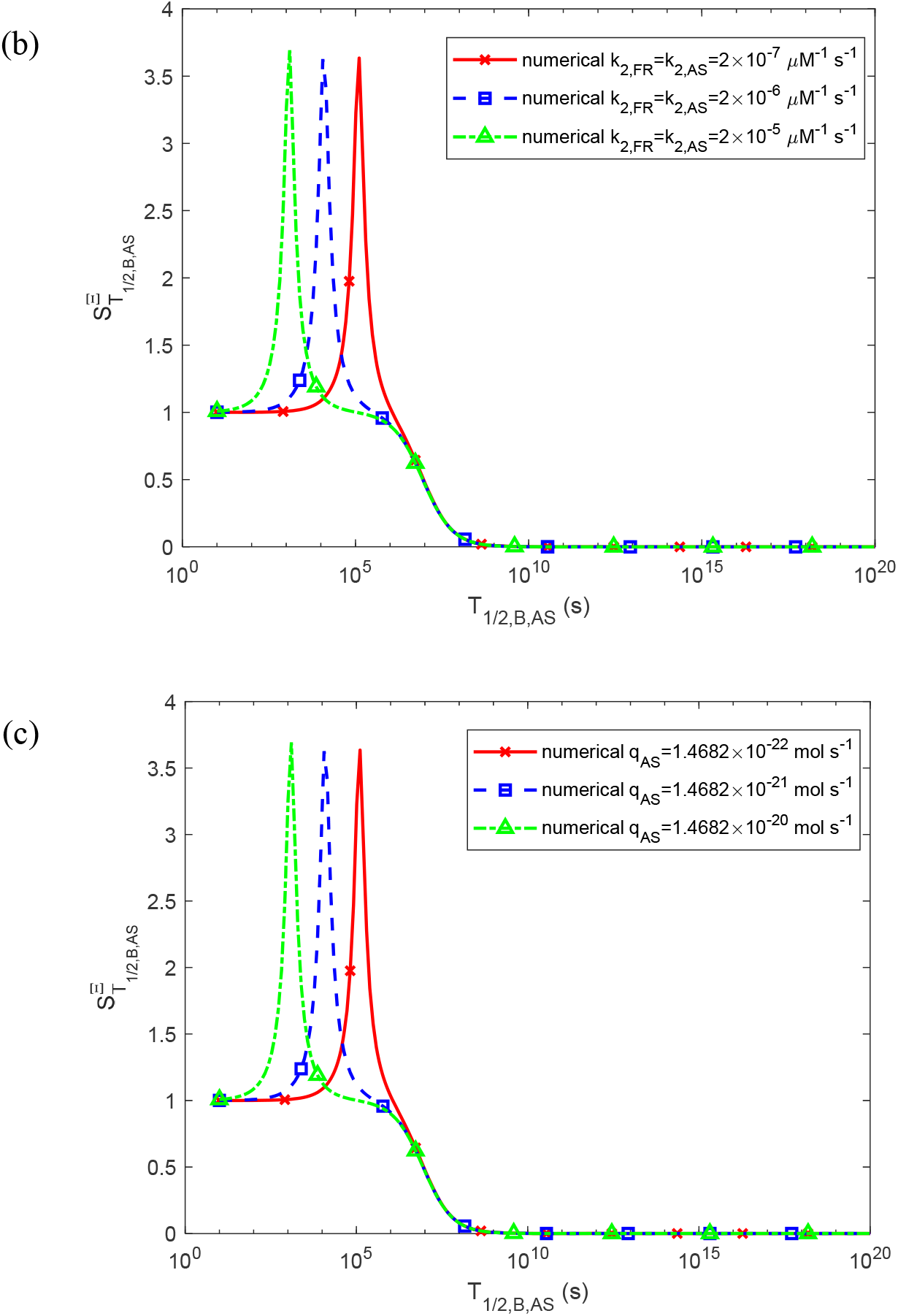
Sensitivity of accumulated toxicity, Ξ, to the half-life of free α-syn aggregates, *T*_1/ 2, *B, AS*_, is examined for three different values of (a) the rate constants that describe the nucleation of membrane fragments and α-syn aggregates, *k*_1,*FR*_ and *k*_1, *AS*_ (*k*_2, *FR*_ = *k*_2, *AS*_ = 2×10^−6^ μM^−1^ s^−1^, *q*_*FR*_ =1.57×10^−28^ mol s^−1^, *q*_*AS*_ = 1.47 ×10^−21^ mol s^−1^), (b) the rate constants that describes the autocatalytic growth of membrane fragments and α-syn aggregates, *k*_2, *FR*_ and *k*_2, *AS*_ (*k*_1,*FR*_ = *k*_1, *AS*_ = 3×10^−7^ s^−1^, *q*_*FR*_ =1.57×10^−28^ mol s^−1^, *q*_*AS*_ = 1.47 ×10^−21^ mol s^−1^), (c) the production rate of α-syn monomers, *q*_*AS*_. (*k*_1,*FR*_ = *k*_1, *AS*_ = 3×10^−7^ s^−1^, *k*_2, *FR*_ = *k*_2, *AS*_ = 2×10^−6^ μM^−1^ s^−1^). *T*_1/ 2, *A, AS*_, *T*_1/ 2, *A, FR*_, and *T* _1/ 2, *B, FR*_ were kept at their values given in Table S2.

